# THE HARD AND SOFT OF IT: THE ROLE OF SUBSTRATE IN PATTERNS OF PHASE DOMINANCE AND PHENOLOGY IN *GRACILARIA VERMICULOPHYLLA*

**DOI:** 10.1101/2025.06.17.660238

**Authors:** Stacy A. Krueger-Hadfield, Alexis P. Oetterer, Darian M. Kelley, Will H. Ryan

**Affiliations:** Department of Biology, University of Alabama at Birmingham, Birmingham, AL, 35294, USA; Virginia Institute of Marine Science Eastern Shore Laboratory, Wachapreague, VA, 23480, USA Alexis P. Oetterer

**Keywords:** Barrier Island System, Chesapeake Bay, Invasion, Life cycle, Marsh, Mudflat, Reproduction, Seaweed

## Abstract

Despite the importance of the seasonal timing of life cycle events for understanding population dynamics, we lack information on the phenology of most macroalgal species. The red macroalga *Gracilaria vermiculophylla* is now common in both hard and soft bottom habitats following its invasion throughout the Northern Hemisphere. In sites with sufficient hard substratum, thalli are fixed by holdfasts to hard substrate. By contrast, in sites with soft substratum, thalli are free-living and either drift free-floating or anchored by the tube building polychaete *Diopatra cuprea*. The difference in substrate type has profound implications on the evolutionary ecology of *G. vermiculophylla* populations whereby the life cycle is disrupted in soft bottom habitats leading to tetrasporophytic dominance. To investigate these patterns across seasons, we determined the reproductive state via microscopy and phase via a sex-linked PCR assay for >4,800 thalli over the course of three years at hard and soft bottom sites with fixed and free-living *G. vermiculophylla* thalli, respectively, along the Eastern Shore of Virginia and Maryland. At hard bottom sites, most thalli were reproductive in the summer and sites were composed of gametophytes and tetrasporophytes, though there was a tetrasporophytic bias. By contrast, soft bottom sites were overwhelming tetrasporophyte dominated and fewer thalli were reproductive at any point in the year. Moreover, each soft bottom site displayed different patterns in the proportion of reproductive thalli, with *G. vermiculophylla* abundance fluctuating through time. Phenology surveys critical for understanding the spread of introduced macroalgae, including the on-going spread of *G. vermiculophylla*.

## INTRODUCTION

Understanding and observing phenology, or the timing of recurring life history events, transcends scientific inquiry as the seasonal progression of these events in plants and animals is something we all perceive, such as tree leafout in spring followed by the loss of those very same leaves in the autumn. The study of phenology has gained increasing attention in response to climate change whereby shifts in the timing of life cycle events affect not only reproduction and survival of individuals of given species but also cascade upwards to significant ramifications for ecosystem structure and function (e.g., Parmesan, 2006; Tang et al., 2016; Walther et al., 2002). The interplay between temperature and photoperiod are important drivers of phenological patterns in angiosperms, such as flowering, though predictions at the community level remain challenging (see Flynn & Wolkovich, 2018 as an example). Similarly, in animals, migratory behavior also varies across species and even ecosystems as different taxa have different drivers inducing migration (reviewed in Shaw, 2016). Understanding the origin of variation in phenology is critical to forecast how species will respond in a changing environment (De Lisle et al., 2022), whereby optima may shift in response to environmental variation across years in seasonally variable habitats (Hadfield, 2016). Therefore, these shifts in the timing of critical life history events are bellwethers of environmental change (Cleland et al., 2007; Inouye, 2022).

Compared to angiosperms in terrestrial ecosystems, our understanding of phenological changes in macroalgae is more limited (reviewed by de Bettignies et al., 2018), despite greater changes in seasonal isotherms in the ocean (Burrows et al., 2012). Studies of macroalgal phenology are often more challenging than similar studies in angiosperms as macroalgae often do not have readily observable reproductive structures, such as flowers. Moreover, macroalgal life cycles are composed of alternations between independent gametophytes and sporophytes that may respond in unique ways to abiotic cues, complicating predictions about how temperature or other abiotic variables may influence phenological patterns (see as an example, the review by Murray & Dixon, 1992). Much of the early work on these patterns, specifically on photoperiod, were undertaken in species with heteromorphic alternations – in which gametophytes and sporophytes can easily be distinguished – and under controlled laboratory conditions (e.g., Dring, 1984). For other heteromorphic gametophytes and sporophytes, phenology studies are challenging when one phase is microscopic, such as the Bangiales, limiting studies (see as an example, Gossard, 2024). Yet, for many macroalgae with isomorphic alternations, we often rely on reproductive material to distinguish phases and sexes, with many studies unable to distinguish between vegetative gametophytes and sporophytes (e.g., Arai et al., 2024; Bhattacharya, 1985; Morley et al., 2003). In their review, de Bettignies et al. (2018) found many phenological studies were undertaken in the red algal order Gigartinales in a heyday of such studies in the 1980s. The focus on the Gigartinales was in part due the ability to distinguish between isomorphic phases when thalli are vegetative using chemical tests (e.g., resorcinol, Lazo et al., 1989). We have mostly focused efforts in the field in hard bottom habitats where gametophytes and sporophytes are fixed to hard substrate by a holdfast (e.g., Dyck & DeWreede, 1995). Largely, these studies have also focused on macroalgal species in temperate environments (de Bettignies et al. 2018, see also review by Olsen et al., 2020 on macroalgal reproductive systems). Thus, there are critical knowledge gaps across macroalgae in their reproduction and the timing of important life history events, rendering it difficult to predict responses to environmental change.

Despite the lack of hard substrate, there are many macroalgae that are found free-living in a variety of soft bottom habitats. Norton and Mathieson (1983) lamented the general dearth of knowledge about the evolutionary ecology of free-living algae and the elapse of 40 years has not greatly improved our understanding. Part of this may be confusion over terminology. Krueger-Hadfield et al. (2025) differentiated among types of free-living algae because secondarily re-attached algae are neither floating nor drifting but were at some point attached by a holdfast before being dislodged and subsequent re-attachment. As a result, different types of free-living macroalgae will have distinct proximate demographic histories influencing their evolutionary ecology. Free-living algae can be found as (i) free-floating and/or drifting thalli, (ii) animal-anchored thalli (used as decoration by polychaetes, crabs, or urchins), or (iii) re-attached thalli via rhizoids, haptera, secondary discs, or some other structure. Yet, Norton and Mathieson (1983) suggested all forms of free-living algae were only capable of vegetative fragmentation and following detachment, would never become reproductive. Unlike this prediction, Krueger-Hadfield et al. (2023) found a peak in the late boreal summer in which reproductive free-living thalli of *Gracilaria vermiculophylla* (Ohmi) Papenfuss were found at a soft bottom site in the non-native range in South Carolina, USA. The onset of reproduction was strongly strong associated with increasing temperatures (Krueger-Hadfield et al., 2023). Moreover, while tetrasporophytes dominated the population throughout the year of monthly surveys, the increase in reproductive, free-living thalli could not have been entirely due to reproductive tetrasporophytes becoming detached from nearby hard bottom populations (Krueger-Hadfield et al., 2023). Thus, there remain many questions about the phenology of free-living algae that require repeated sampling through time to document their phenological calendar of life history events from which we can gain a better understanding of their evolutionary ecology.

Here, we continue work on the phenological patterns in *Gracilaria vermiculophylla*, a red alga now found throughout the Northern Hemisphere following its invasion from the Northwest Pacific (Krueger-Hadfield et al., 2017). While previous work has focused on native versus non-native thalli to understand more about this prolific invasion (e.g., Hammann et al., 2016; Murren et al., 2022; Sotka et al., 2018), almost all native populations were sampled in hard bottom habitats (e.g., rocky shores or mudflats with abundant hard substrate) whereas a majority of non-native populations were sampled in soft bottom habitats (e.g., mudflats with no hard substrate). Krueger-Hadfield et al. (2016) described patterns of sexual reproduction and the occurrence of gametophytes and tetrasporophytes in the native range as compared to the dominance of asexual reproduction and tetrasporophytes in the non-native range. However, this difference in the prevailing reproductive mode and the phase ratio is unlikely to reflect a simple native versus non-native dichotomy. Our current knowledge about the reproductive system and phase ratio in *G. vermiculophylla* has been influenced by a bias in the choice of sampling sites, as subsequent surveys have revealed thalli in both hard and soft bottom habitats in the native and non-native ranges (Krueger-Hadfield et al., 2017; 2018). The disruption of the life cycle in which free-living tetrasporophytes dominate and asexual reproduction prevails is driven largely by the lack of substrate in soft bottom habitats and some of these patterns are further magnified by the invasion (e.g., Baker’s Law in which reproductive mode flexibility enhances colonization success, Baker, 1955; Pannell et al., 2015). Yet, we require more detailed within and among site sampling in both the native and non-native range to understand invasion dynamics across spatial scales in haploid-diploid taxa (see Krueger-Hadfield, 2020; Krueger-Hadfield & Hoban, 2016).

The first steps are baseline surveys and the documentation of phenological patterns in both hard and soft bottom sites. Currently, the handful of studies in *Gracilaria vermiculophylla* populations are confounded by the invasion and habitat type. In the native range, phenological studies have taken place in hard bottom habitats. Terada et al. (2000) documented a summer peak in reproductive thalli, the majority of which were tetrasporophytes, at a site in Hokkaido (northern Japan, ∼41°N). However, most sampled thalli were vegetative, in which the phase and sex could not be determined. At a site on Kyushu Island (∼33°N) in southern Japan, reproductive tetrasporophytes were present year-round and dominated the population (Muangmai et al., 2014). In the non-native range, Abreu et al. (2011) also found reproductive thalli throughout the year at a site in Portugal but never observed male gametophytes. Many thalli were vegetative at the time of collection and the phase was not determined. This site in Portugal had sufficient hard substrate and therefore differed from the site studied by Krueger-Hadfield et al. (2023) in which there was little hard substrate. Unlike Abreu et al. (2011), Krueger-Hadfield et al. (2023) were able to determine the phase of all thalli collected throughout the year of monthly samples, observing mostly tetrasporophytes and a peak in reproduction in late summer. As the *G. vermiculophylla* populations studied in Portugal (Abreu et al., 2011) and South Carolina (Krueger-Hadfield et al., 2023) are the result of distinct invasions (Krueger-Hadfield et al., 2017), we cannot tease apart the differences in phenological patterns that may arise from the invasion or substrate type. We further cannot compare patterns of phenology confounded by the introduction of *G. vermiculophylla* throughout the Northern Hemisphere as the invasion history may influence demography (e.g., Sakai et al., 2001) and phenology (e.g., Ito et al., 2021).

To begin to establish baseline patterns with which we can delve more deeply into the evolutionary ecology of haploid-diploid life cycles, we surveyed hard and soft bottom habitats colonized by *Gracilaria vermiculophylla* is now found along the coast of the Eastern Shore of Virginia and Maryland. This region falls in the center of the species’ latitudinal range on this coast and has been colonized since at least the mid-1900s (Krueger-Hadfield et al., 2017). Previous sampling has demonstrated that these sites are genetically similar (Krueger-Hadfield et al., 2017) and established the occurrence of gametophytes and tetrasporophytes in hard bottom sites and tetrasporophytes in soft bottom sites (Krueger-Hadfield et al., 2016; 2017; Krueger-Hadfield & Ryan, 2020). We sampled *G. vermiculophylla* thalli at three hard bottom and two soft bottom sites once per boreal season from 2020 to 2023 (an additional soft bottom site was added in the summer of 2022). We predicted strong seasonal periodicity in reproduction with a peak in the summer in both hard and soft bottom sites based on previous phenology studies (Krueger-Hadfield et al., 2023; Terada et al., 2000) and prior observations (S.A. Krueger-Hadfield, *personal observations*). In hard bottom sites, we expected to observe a slight tetrasporophytic bias as is common in the Gracilariales (Kain & Destombe, 1995) and 1:1 sex ratio in the gametophytes as expected from Mendelian segregation of the sex determining region (Krueger-Hadfield et al., 2023; Lipinska et al., 2024). In soft bottom sites, we hypothesized most if not all thalli collected would be tetrasporophytes (see Krueger-Hadfield et al., 2016). Moreover, we predicted we would find reproductive tetrasporophytes as previously found by Krueger-Hadfield et al. (2023). This is one of the first studies of which we are aware in which the phenology and the determination of phase for each thallus has been described in the same species with both fixed and free-living thalli.

## MATERIALS AND METHODS

### Algal collections and identification

We sampled *Gracilaria vermiculophylla* thalli at five sites along the Eastern Shore of Maryland and Virginia from 2020 to 2023, with an additional sixth site added in summer of 2022 (Figure 1, Table 1, Table S1). We collected fixed thalli haphazardly at the three hard bottom sites – AHC, Ape Hole Creek; CCB, Cape Charles Beach; and MAG, Cushman’s Landing at the end of Magotha Road. We individually bagged thalli, with each thallus kept in numerical spatial sampling order using scroll bags (Krueger-Hadfield, 2021) before processing immediately following collection at the Virginia Institute of Marine Science Eastern Shore Laboratory (VIMS ESL). We subsampled upright thalli from distinct holdfasts to avoid sampling the same genet (i.e., genotype) twice either in the field or in the laboratory.

**Figure 1.**
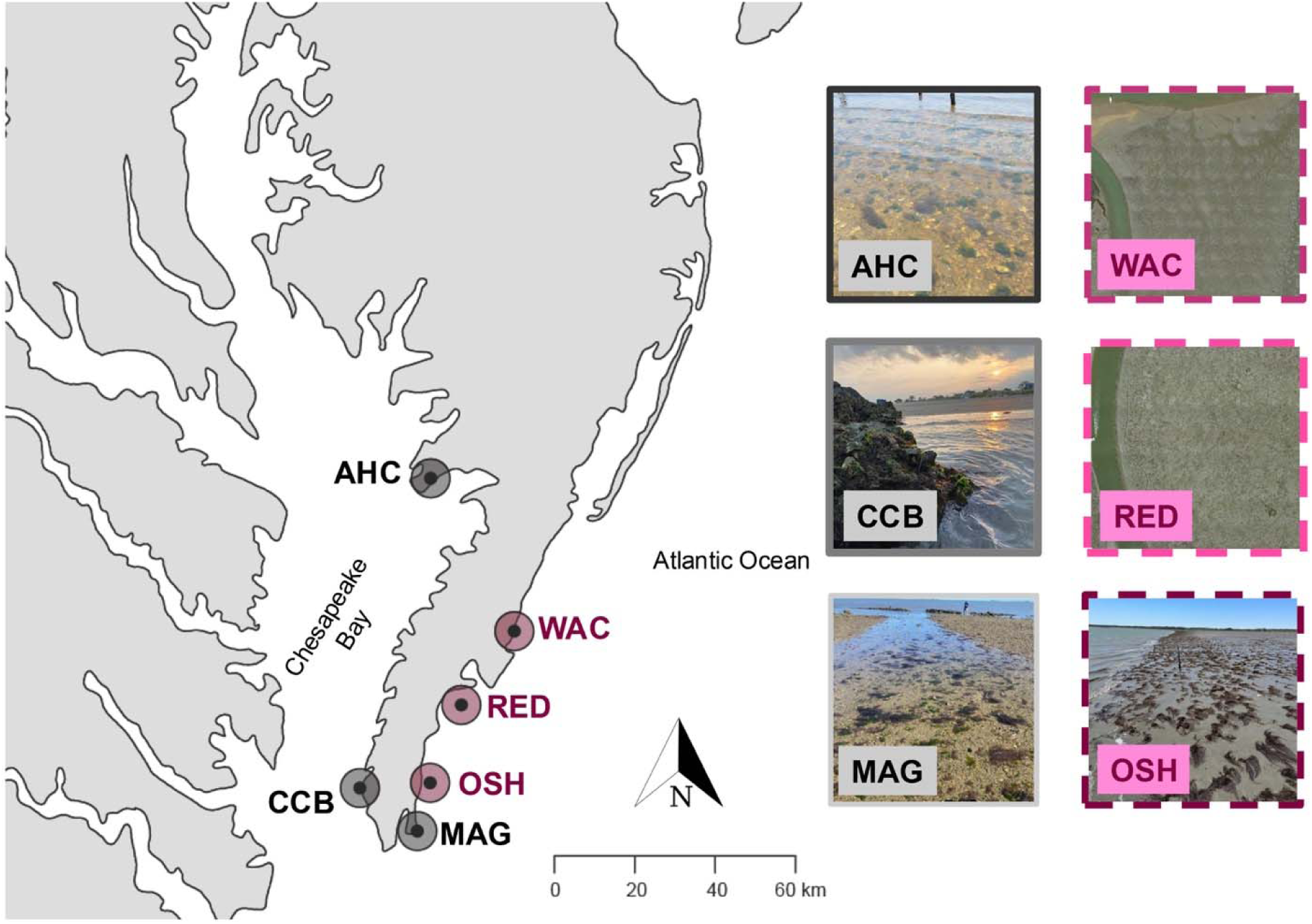
Map of the Delmarva Peninsula showing the six sites sampled to document the phenology of *Gracilaria vermiculophylla* populations in hard and soft bottom habitats. AHC, Ape Hole Creek; CCB, Cape Charles Beach; and MAG, Magotha are hard bottom sites (shown in shades of black/gray). OSH, Hillcrest; RED, Fowling Point; and WAC, Upper Haul Over are soft bottom sites (shown in shades of maroon). Detailed site information is provided in Table 1. Site photo credits: AHC, CCB, MAG, OSH, S.A. Krueger-Hadfield; WAC, RED, PG Ross.

**Table 1.** Metadata for each site in which *Gracilaria vermiculophylla* thalli were sampled along the Delmarva Peninsula. Site, Site abbreviation and name; Habitat type, hard bottom (and type) or soft bottom; Latitude; Longitude; Season, boreal summer, boreal fall, boreal winter, or boreal spring; Collection date; Salinity, in PSU; N*_Diopatra_*, the average number of *Diopatra cuprea* tube caps per m^2^; N, sample size of thalli; P(Tetra), proportion of sample that was a tetrasporophyte; P(Gameto), proportion of sample that was a gametophyte; P(Fem), proportion of sample that was a female gametophyte; P(Male), proportion of sample that was a male gametophyte; P_HD_, ploidy diversity; P_HD_, sex diversity. *P_HD_*and *P_FM_* describe the phase and sex diversity, respectively, at a given sampling site and collection date is the ploidy diversity metric that describes the amount of life cycle stage diversity at each sampling point (Krueger-Hadfield et al. 2019). ** denotes *p* < 0.05 and * denotes 0.05 < *p* < 0.1 for either *P_HD_*and *P_FM_* values that differ from √2:1 or 1:1, respectively, following deviations from a binomial distribution. Gametophytes were rare at all soft bottom sites, therefore, deviations from a 1:1 sex ratio were not tested at these sites.

| Site | Habitat Type | Latitude | Longitude | Season | Collection date | Salinity | $N_{Diopatra}$ | N | P(Tetra) | P(Gameto) | P(Fem) | P(Male) | $P_{HD}$ | $P_{FM}$ |
| --- | --- | --- | --- | --- | --- | --- | --- | --- | --- | --- | --- | --- | --- | --- |
| AHC – | Hard bottom | 37.95843758 | -75.82432691 | Summer | 2 Jul 2020 | 16 | - | 105 | 0.50 | 0.50 | 0.27 | 0.23 | 0.86** | 0.94 |
| Ape | – shell hash |  |  | Fall | 16 Oct 2020 | 20 | - | 107 | 0.68 | 0.32 | 0.21 | 0.11 | 0.54** | 0.71** |
| Hole |  |  |  | Winter | 28 Jan 2021 | 17 | - | 81 | 0.57 | 0.43 | 0.16 | 0.27 | 0.73** | 0.74** |
| Creek |  |  |  | Spring | 25 Apr 2021 | 20 | - | 100 | 0.59 | 0.41 | 0.20 | 0.21 | 0.69** | 0.98 |
|  |  |  |  | Summer | 18 Jul 2021 | 16 | - | 100 | 0.59 | 0.41 | 0.31 | 0.10 | 0.69** | 0.49** |
|  |  |  |  | Fall | 3 Oct 2021 | 15 | - | 102 | 0.67 | 0.33 | 0.12 | 0.22 | 0.56** | 0.71** |
|  |  |  |  | Winter | 2 Feb 2022 | 17 | - | 51 | 0.63 | 0.37 | 0.22 | 0.16 | 0.63** | 0.84 |
|  |  |  |  | Spring | 18 Apr 2022 | 15 | - | 50 | 0.64 | 0.36 | 0.18 | 0.18 | 0.61** | 1.00 |
|  |  |  |  | Summer | 19 Jul 2022 | 15 | - | 98 | 0.60 | 0.40 | 0.22 | 0.17 | 0.67** | 0.87* |
|  |  |  |  | Fall | 19 Oct 2022 | 21 | - | 51 | 0.71 | 0.29 | 0.18 | 0.12 | 0.50** | 0.80 |
|  |  |  |  | Winter | 23 Feb 2023 | 17 | - | 97 | 0.55 | 0.45 | 0.19 | 0.27 | 0.77** | 0.82* |
|  |  |  |  | Spring | 20 Apr 2023 | 17 | - | 101 | 0.58 | 0.42 | 0.24 | 0.18 | 0.70** | 0.86* |
|  |  |  |  | Summer | 18 Jul 2023 | 19 | - | 114 | 0.48 | 0.52 | 0.31 | 0.21 | 0.88** | 0.81** |
| CCB – | Hard bottom - | 37.2715101 | -76.02319154 | Summer | 1 Jul 2020 | 20 | - | 49 | 0.53 | 0.47 | 0.27 | 0.20 | 0.80** | 0.87 |
| Cape | jetty |  |  | Fall | 17 Oct 2020 | 24 | - | 100 | 0.57 | 0.43 | 0.15 | 0.28 | 0.73** | 0.70** |
| Charles |  |  |  | Winter | 30 Jan 2021 | 25 | - | 99 | 0.39 | 0.60 | 0.21 | 0.39 | 0.96* | 0.70** |
| Beach |  |  |  | Spring | 28 Apr 2021 | 19 | - | 50 | 0.60 | 0.40 | 0.12 | 0.28 | 0.68** | 0.60** |
|  |  |  |  | Summer | 26 Jul 2021 | 20 | - | 97 | 0.59 | 0.41 | 0.30 | 0.11 | 0.70** | 0.55** |
|  |  |  |  | Fall | 2 Oct 2021 | 25 | - | 50 | 0.80 | 0.20 | 0.14 | 0.06 | 0.34** | 0.60** |
|  |  |  |  | Winter | 28 Jan 2022 | 26 | - | 100 | 0.42 | 0.58 | 0.21 | 0.37 | 0.98* | 0.72** |
|  |  |  |  | Spring | 15 Apr 2022 | 19 | - | 100 | 0.46 | 0.54 | 0.28 | 0.26 | 0.92** | 0.96 |
|  |  |  |  | Summer | 18 Jul 2022 | 26 | - | 96 | 0.68 | 0.32 | 0.11 | 0.21 | 0.55** | 0.71** |
|  |  |  |  | Fall | 21 Oct 2022 | 25 | - | 48 | 0.60 | 0.40 | 0.23 | 0.17 | 0.67** | 0.84 |
|  |  |  |  | Winter | 19 Feb 2023 | 24 | - | 109 | 0.50 | 0.50 | 0.28 | 0.23 | 0.86** | 0.91* |
|  |  |  |  | Spring | 15 Apr 2023 | 26 | - | 100 | 0.54 | 0.45 | 0.19 | 0.26 | 0.78** | 0.84* |
|  |  |  |  | Summer | 17 Jul 2023 | 20 | - | 99 | 0.57 | 0.43 | 0.25 | 0.18 | 0.74** | 0.84* |
| MAG – | Hard bottom | 37.17602001 | -75.94173218 | Summer | 1 Jul 2020 | 29 | - | 126 | 0.52 | 0.48 | 0.25 | 0.22 | 0.81** | 0.93* |
| Magotha | – shell hash |  |  | Fall | 17 Oct 2020 | 31 | - | 104 | 0.62 | 0.38 | 0.23 | 0.14 | 0.64** | 0.77** |
| Road |  |  |  | Winter | 30 Jan 2021 | 30 | - | 98 | 0.55 | 0.45 | 0.10 | 0.35 | 0.76** | 0.45** |
|  |  |  |  | Spring | 24 Apr 2021 | 31 | - | 100 | 0.60 | 0.40 | 0.23 | 0.17 | 0.68** | 0.85* |
|  |  |  |  | Summer | 17 Jul 2021 | 30 | - | 101 | 0.67 | 0.33 | 0.24 | 0.09 | 0.55** | 0.54** |
|  |  |  |  | Fall | 2 Oct 2021 | 35 | - | 102 | 0.60 | 0.40 | 0.17 | 0.23 | 0.68** | 0.83* |
|  |  |  |  | Winter | 28 Jan 2022 | 30 | - | 104 | 0.58 | 0.42 | 0.18 | 0.24 | 0.72** | 0.86* |
|  |  |  |  | Spring | 15 Apr 2022 | 27 | - | 98 | 0.72 | 0.28 | 0.09 | 0.18 | 0.46** | 0.67** |
|  |  |  |  | Summer | 17 Jul 2022 | 27 | - | 73 | 0.57 | 0.42 | 0.22 | 0.21 | 0.72** | 0.97 |
|  |  |  |  | Fall | 21 Oct 2022 | 33 | - | 50 | 0.58 | 0.42 | 0.14 | 0.28 | 0.71** | 0.67* |
|  |  |  |  | Winter | 19 Feb 2023 | 29 | - | 101 | 0.53 | 0.47 | 0.31 | 0.16 | 0.79** | 0.68** |
|  |  |  |  | Spring | 15 Apr 2023 | 34 | - | 147 | 0.61 | 0.39 | 0.15 | 0.24 | 0.66** | 0.77** |
|  |  |  |  | Summer | 16 Jul 2023 | 30 | - | 126 | 0.49 | 0.51 | 0.31 | 0.20 | 0.86** | 0.78** |
| OSH – | Soft bottom – | 37.28309983 | -75.90591928 | Summer | 15 Jul 2022 | 25 | 32.5 | 43 | 1.0 | 0.00 | 0.00 | 0.00 | 0.00** | - |
| Hillcrest | free-living or |  |  | Fall | 25 Oct 2022 | 34 | 24.6 | 31 | 0.97 | 0.03 | 0.03 | 0.0 | 0.05** | - |
|  | worm |  |  | Winter | 20 Feb 2023 | 33 | 36.6 | 29 | 1.0 | 0.00 | 0.00 | 0.00 | 0.00** | - |
|  | anchored |  |  | Spring | 18 Apr 2023 | 36 | 34.3 | 26 | 1.0 | 0.00 | 0.00 | 0.00 | 0.00** | - |
|  |  |  |  | Summer | 14 Jul 2023 | 34 | 29.7 | 56 | 1.0 | 0.00 | 0.00 | 0.00 | 0.00** | - |
| RED – | Soft bottom – | 37.45585769 | -75.8183666 | Summer | 28 Jun 2020 | 28 | - | 61 | 0.92 | 0.08 | 0.07 | 0.01 | 0.14** | - |
| Fowling | free-living or |  |  | Fall | 14 Oct 2020 | 24 | - | 38 | 0.97 | 0.03 | 0.00 | 0.03 | 0.04** | - |
| Point | worm |  |  | Winter | 29 Jan 2021 | 34 | - | 30 | 1.0 | 0.00 | 0.00 | 0.00 | 0.00** | - |
|  | anchored |  |  | Spring | 27 Apr 2021 | 28 | - | 52 | 0.96 | 0.04 | 0.00 | 0.04 | 0.07** | - |
|  |  |  |  | Summer | 22 Jul 2021 | 29 | - | 65 | 0.92 | 0.08 | 0.03 | 0.05 | 0.13** | - |
|  |  |  |  | Fall | 1 Oct 2021 | 37 | - | 38 | 1.0 | 0.00 | 0.00 | 0.00 | 0.00** | - |
|  |  |  |  | Winter | 31 Jan 2022 | 35 | - | 30 | 0.93 | 0.07 | 0.00 | 0.07 | 0.12** | - |
|  |  |  |  | Spring | 22 Apr 2022 | 31 | - | 34 | 0.94 | 0.06 | 0.06 | 0.00 | 0.10** | - |
|  |  |  |  | Summer | 14 Jul 2022 | 25 | 22.7 | 41 | 0.95 | 0.05 | 0.00 | 0.05 | 0.08** | - |
|  |  |  |  | Fall | 26 Oct 2022 | 33 | 13.7 | 13 | 1.0 | 0.00 | 0.00 | 0.00 | 0.00** | - |
|  |  |  |  | Winter | 21 Feb 2023 | 35 | 20.6 | 23 | 1.0 | 0.00 | 0.00 | 0.00 | 0.00** | - |
|  |  |  |  | Spring | 19 Apr 2023 | 36 | 17.7 | No thalli present<sup>1</sup> |  |  |  |  |  |  |
|  |  |  |  | Summer | 17 Jul 2023 | 30 | 21.7 | 23 | 0.87 | 0.13 | 0.13 | 0.00 | 0.22** | - |
| WAC – | Soft bottom – | 37.61969909 | -75.66938578 | Summer | 6 Jul 2020 | 30 | - | 74 | 1.0 | 0.00 | 0.00 | 0.00 | 0.00** | - |
| Upper | free-living or |  |  | Fall | 15 Oct 2020 | 29 | - | 43 | 0.98 | 0.02 | 0.02 | 0.00 | 0.04** | - |
| Haul | worm |  |  | Winter | 27 Jan 2021 | 32 | - | 65 | 0.98 | 0.02 | 0.02 | 0.00 | 0.03** | - |
| Over | anchored |  |  | Spring | 26 Apr 2021 | 35 | - | 68 | 0.94 | 0.06 | 0.00 | 0.06 | 0.10** | - |
|  |  |  |  | Summer | 23 Jul 2021 | 31 | - | 93 | 1.0 | 0.00 | 0.00 | 0.00 | 0.00** | - |
|  |  |  |  | Fall | 6 Oct 2021 | 36 | - | 66 | 1.0 | 0.00 | 0.00 | 0.00 | 0.00** | - |
|  |  |  |  | Winter | 1 Feb 2022 | 34 | - | 54 | 1.0 | 0.00 | 0.00 | 0.00 | 0.00** | - |
|  |  |  |  | Spring | 19 Apr 2022 | 31 | - | 62 | 0.98 | 0.02 | 0.02 | 0.00 | 0.03** | - |
|  |  |  |  | Summer | 18 Jul 2022 | 27 | 34.3 | 37 | 1.0 | 0.00 | 0.00 | 0.00 | 0.00** | - |
|  |  |  |  | Fall | 24 Oct 2022<sup>2</sup> | 32 | 24.6 | 20 | 0.80 | 0.20 | 0.00 | 0.20 | 0.34** | - |
|  |  |  |  | Winter | 22 Feb 2023 | 36 | 33.1 | 23 | 0.91 | 0.09 | 0.00 | 0.09 | 0.15** | - |
|  |  |  |  | Spring | 17 Apr 2023 | 34 | 22.3 | 11 | 1.0 | 0.00 | 0.00 | 0.00 | 0.00** | - |
|  |  |  |  | Summer | 18 Jul 2023 | 30 | 25.7 | 22 | 0.95 | 0.05 | 0.05 | 0.00 | 0.08** | - |

<sup>1</sup> No *G. vermiculophylla* thalli were present at Fowling Point or nearby sites on this collection date.

At soft bottom sites – OSH, Hillcrest out of Oyster Harbor; RED, Fowling Point out of Willis Wharf; and WAC, Upper Haul Over, Burton’s Bay out of Wachapreague (Figure 1, Table 1, Table S1) – free-living *G. vermiculophylla* thalli are either drifting or anchored to the tube caps of the tube-building polychaete *Diopatra cuprea*. Instead of haphazard sampling, at soft bottom sites, we sampled all thalli present in each of seven 0.25 m^2^ quadrats along a 30-m transect. These quadrats were sampled in the same area of the site at each collection event. All thalli from a quadrat were placed into 5-gallon paint strainer mesh bags. In the laboratory, a single piece of contiguous thallus was disentangled and used for downstream analyses. From summer 2022 onwards, in each quadrat, we also recorded the number of *Diopatra cuprea* tube caps, assuming the number of tube caps represents the number of polychaetes, to better monitor the *G. vermiculophylla* biomass.

We determined reproductive state using a dissecting microscope (40x) to distinguish reproductive tetrasporophytes, reproductive male gametophytes, and cystocarpic female gametophytes (see Krueger-Hadfield et al., 2018). If a thallus had no visible reproductive structures, it was considered vegetative. We preserved a piece of each thallus in silica gel for subsequent DNA extraction. Any visible cystocarps were excised prior to preservation in silica gel to remove the introduction of exogenous DNA from paternal contributions to the carposporophyte.

We considered a female gametophyte as ‘reproductive’ when the thallus bore cystocarps at the time of collection. This is a conservative estimate of reproductive females as some vegetative thalli may have had mature carpogonia not yet fertilized to form the carposporophyte. This would not have been visible at the time of collection and the thallus would be considered vegetative.

### Abiotic data

We deployed HOBO Pendant Temperature/Light 64K Data Loggers (Cat No. UA-002-64, ONSET, Bourne, MA, USA) at AHC, MAG, OSH, RED, and WAC. We deployed a temperature logger at the Cape Charles Marina (CHR, 37.26482, −76.01854) rather than CCB due to logistical constraints of permanently attaching a logger on the jetties at CCB. The marina is approximately 1.4 km away from CCB. We recorded measurements every ten minutes and exchanged loggers at each collection event. Some loggers were either lost or did not record data for the entire deployment resulting in the data gaps.

We measured salinity with a refractometer at each site and each collection date for point estimates of salinity (PPT).

Continuously recording, fixed-sensor water quality monitoring stations with YSI EXO2 multiparameter sondes (YSI, Yellow Springs, OH) occur at three sites along the Eastern Shore of Virginia run by the VIMS ESL with detailed methods provided in Kelley (2025). We obtained temperature (°C) and salinity (PSU) data from the Wachapreague (37.607692, −75.685809) and Willis Wharf stations (37.511967, −75.806165). Data from Wachapreague were unavailable in 2020 due to COVID-19 associated technical issues, therefore, we used data from 2021 to 2023. Data for the Willis Wharf station are available from 2020 to 2023. There are some months missing throughout this period at each station when equipment was not operational. Additional quality control parameters are reported in Kelley (2025). The Wachapreague station is close to the WAC site and the Willis Wharf station is close to the RED site.

### DNA extraction, PCR, and sex-linked marker assay

As many thalli were vegetative at the time of sampling, we used the sex-linked genetic markers developed by Krueger-Hadfield et al. (2021) and modified in Krueger-Hadfield et al. (2023) to determine phase (and for gametophytes, sex) for all thalli sampled in the summers of 2020, 2021, and 2022, and all vegetative thalli. We used these sex-linked markers on all thalli, including those for which we identified reproductive structures. We adapted a Chelex DNA extraction protocol from Simon et al. (2020) (see also Krueger-Hadfield et al., 2023). We placed ∼1 cm of each silica gel-dried thallus into a Fisherbrand^TM^ 96-well non-skirted PCR plate and then added 200 µL of pre-heated (95 °C) 5% Chelex solution (Bio-Rad Laboratories, Hercules, CA). Each plate was vortexed for 30 seconds, then incubated at 95 °C for 15 minutes with a brief vortex every 5 minutes. Each plate was centrifuged for 3 minutes at 5,600 x *g* after which 180 µL of the supernatant was transferred to a new 96-well plate. Each plate was centrifuged again for 3 minutes at 5,600 x *g* and 150 µL of the supernatant was transferred to a final 96-well plate. For each plate, we had two female gametophytic, two male gametophytic, and two tetrasporophytic positive controls, one negative DNA extraction control, and one negative PCR control.

We used the same PCR assay as Krueger-Hadfield et al. (2023) and amplified the fem_o03 and mal_n09 loci in a 10 µL duplex PCR containing 2 µL of DNA template, 0.5 U GoTAQ Flexi-DNA Polymerase (Promega, Madison, WI), 1x green reaction buffer, 100 µM of each dNTP, 1.5 mM of MgCl_2_, 1 mg/mL bovine serum albumin (BSA), 250 nM female forward and reverse primers, and 350 nM male forward and reverse primers, and the cycling profile: 95 °C for 10 minutes, 35 cycles of 95 °C for 30 s, 59 °C for 30 s, and 72 °C for 30 s, followed by a final extension of 72 °C for 5 min. We visualized 3 µL of each PCR product on a 1.5% agarose gel stained with 6 µL GelRed (Biotium, Fremont, CA). We scored each thallus blind and then matched the phase and sex when applicable based on the sex-linked marker (female: 73 base pairs (bp); male: 270 bp; tetrasporophyte: both 73 bp and 270 bp) with the reproductive state identified at the time of collection.

### Phase ratio, ploidy diversity P_HD_, and sex diversity P_FM_

Using the binomial distribution, we tested whether the gametophyte to tetrasporophyte ratio differed from the predicted ratio of √2:1 (Destombe et al., 1989; Thornber & Gaines, 2004). We chose this prediction of phase ratio to compare the current findings with previous work at several of the same sites (e.g., Krueger-Hadfield et al., 2016) as well as a recent study by Krueger-Hadfield et al. (2023).

We also calculated ploidy diversity (*P_HD_*, Krueger-Hadfield et al., 2019). Since the proportion of tetrasporophytes was greater than 0.41 at all sampling periods, we used 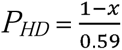 where x is the proportion of tetrasporophytes at a site. When *P_HD_* = 1, the ratio of gametophytes to tetrasporophytes is the predicted √2:1, but when *P_HD_* = 0, there is only one life cycle stage – gametophyte or sporophyte. In our study, as *P_HD_* approaches 0, there are only tetrasporophytes present at a site.

For all the hard bottom sites with fixed thalli, we also calculated sex diversity (*P_FM_*, Krueger-Hadfield et al., 2019). Analogous to ploidy diversity, maximum diversity occurs at a 1:1 sex ratio (i.e., 50% female and 50% male). We used the equation 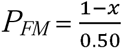, where *x* is the proportion of females or males depending on the observed ratios.

### Analyses to assess within and among site variation

All analyses were conducted in R ver. 4.3.3. (R Core Team, 2024).

We compared the *Diopatra cuprea* tube cap counts across the soft bottom sites (OSH, RED, and WAC) from summer 2022 to summer 2023 using a generalized linear mixed model (GLMM) with a negative binomial distribution and a log link function in the glmmTMB package (McGillycuddy et al., 2025). The model included site and season as fixed effects, and quadrat as a random intercept to account for spatial replication. Pairwise comparisons were performed using the emmeans package (Lenth, 2025) with Tukey adjustment for multiple tests.

We investigated several different components of the *Gracilaria vermiculophylla* phenological calendar using a beta regression model with a logit link function in the glmmTMB package to test the effects of: (i) site and season on ploidy diversity (*P_HD_*), (ii) hard bottom site and season on sex diversity (*P_FM_*), (iii) site and season on the proportion of reproductive thalli, and (iv) site and the mean temperature (°C) 30 days before thallus collection on the proportion of reproductive thalli. First, we analyzed the difference among sites on *P_HD_*. While *P_HD_* was normally distributed in hard bottom sites, it was strongly right skewed in the soft bottom sites, leading to a bimodal distribution when hard and soft bottoms sites were combined. We considered season (Winter, Spring, Summer, Fall; Fall as the reference level) and site (AHC, CCB, MAG, OSH, RED, and WAC; AHC as the reference level) as fixed factors. We treated year (sampling years from 2020 to 2022 or 2023) as a random factor. We adjusted the *P_HD_* values by 1×10^−6^, ensuring the values were between 0 and 1, but not equal to 0 or 1. We compared seasons using contrasts in the emmeans package (Lenth, 2025) with Tukey adjustment for multiple tests.

Second, in hard bottom sites, we explored how the fixed effects of site (AHC, CCB, MAG; AHC as the reference level) and season (Winter, Spring, Summer, Fall; Fall as the reference level) with the random effect of year (sampling years from 2020 to 2022) on *P_FM_*. While we found the occasional gametophyte in soft bottom sites, the sample sizes were too low to perform any robust analysis. Thus, in soft bottom sites, we did not calculate *P_FM_*. We adjusted the *P_HD_* values by 1×10^−6^, ensuring the values were between 0 and 1, but not equal to 0 or 1. We compared seasons using contrasts in the emmeans package with Tukey adjustment for multiple tests.

Third, at hard bottom sites, we analyzed the effect of site and season on the proportion of reproductive thalli. We note that reproductive thalli will include cystocarpic females, reproductive males, and reproductive tetrasporophytes, whereas at soft bottom sites, the proportion of reproductive thalli will likely only include tetrasporophytes (see Krueger-Hadfield et al., 2016). In case there are differences in reproductive periodicity between gametophytes and tetrasporophytes, we first compared the distribution of the proportion of reproductive thalli to the proportion of reproductive tetrasporophytes using a Wilcoxon rank sum test. Since there were no differences between the distributions of these two groups (W = 2556.5, *p* = 0.454), we then explored how the fixed effects of site (AHC, CCB, MAG, OSH, RED, WAC; AHC as the reference level) and season (Winter, Spring, Summer, Fall; Fall as the reference level) with the random effect of year (sampling years from 2020 to 2022 and 2023 for OSH) on the proportion of reproductive thalli. We adjusted the proportion by 1×10^−6^, ensuring the values were between 0 and 1, but not equal to 0 or 1. We compared seasons and sites using contrasts in the emmeans package with Tukey adjustment for multiple tests.

Fourth, to investigate the relationship between temperature and the proportion of reproductive thalli, we used temperature data from the Willis Wharf water quality station as that station was the only one from which had uninterrupted and continuous measurements from 2020 to 2023. We then averaged the temperatures over the 30 days prior to each sampling event at each of our six sites. We explored how the fixed effects of site (AHC, CCB, MAG, OSH, RED, WAC; AHC as the reference level) and mean temperature 30 days prior to collection on the proportion of reproductive thalli. We adjusted the proportion by 1×10^−6^, ensuring the values were between 0 and 1, but not equal to 0 or 1. We compared seasons and sites using contrasts in the emmeans package Tukey adjustment for multiple tests.

To assess the temporal relationship between the proportion of cystocarpic female gametophytes and reproductive male gametophytes across seasons, we used a cross-correlation function in base R to evaluate the strength and direction of the correlation using a maximum lag of ± 2 seasons. We chose 2 seasons which is equivalent to approximately half a year to represent a biologically meaningful lag in which males might exhibit reproductive structures before we can observe the development of carposporophytes (e.g., Dyck & DeWreede, 1995). To test the significance of observed cross-correlations between cystocarpic females and reproductive males, we implemented a non-parametric permutation test (N = 1000 permutations). We only performed this analyses in hard bottom sites.

We calculated the change in *P_HD_* and proportion of reproductive thalli between sampling events and used a Spearman rank correlation to test the hypothesis that there is a relationship between *P_HD_* and the proportion of reproductive thalli. As we only sampled thalli at OSH from summer 2022 until summer 2023, we did not include this soft bottom site in these analyses.

### Data visualization

Data organization and figures were prepared using the following packages in R using the packages doBy (Højsgaard & Halekoh, 2023), ggplot2 (Wickham, 2016), cowplot (Wilke, 2024), reshape2 (Wickham, 2007), dplyr (Wickham et al., 2023), and openxlsx (Shauberger & Walker, 2023).

## RESULTS

### Temperature and salinity variation at hard and soft bottom sites

The HOBO loggers at MAG, OSH, RED, and WAC recorded higher maximum temperatures (43.5°C to 49.5°C) than at AHC and CHR (35.0°C and 39.5°C, respectively, Figure S1, Table S2a). The loggers at the two bayside sites – AHC and CHR – were consistently submerged, even at low tide, and therefore were recording water temperature. By contrast, the loggers at the seaside sites at MAG, OSH, RED, and WAC were submerged at high tide and emerged at low tide, measuring water and air temperature, respectively, depending on the tidal cycle. Over the course of the phenology survey, the mean temperature ranged from 16.1°C at MAG to 17.4°C at RED, the lowest temperature was observed at MAG (−9.7°C; range of minimum temperatures: −9.7°C to −4.3°C at CHR), and the maximum temperatures ranged from 35.0°C at AHC to 49.5°C at OSH.

Salinity ranged from 15 PPT to 37 PPT over the course of the study from our point estimates on collection dates at each site (Table 1). At AHC, salinity was always below 20 PPT at all sampling dates, whereas at CCB, the other bayside site, salinity ranged from 19 – 26 PPT. All seaside sites, salinity was greater than 24 PPT.

### Temperature and salinity variation at Willis Wharf and Wachapreague

At the water quality stations at Wachapreague (Table S2b) and Willis Wharf (Table 2c), a floating pump delivers a continuous water sample to a chamber where the sonde is located on land. The pump is floating, and the samples are collected at a fixed depth ∼50 cm below the surface, regardless of tidal height. Data was recorded for both salinity and temperature every 15 minutes (Figure S2). At Willis Wharf, from 2020 to 2023, the lowest temperature recorded in Parting Creek was 0.003°C and the highest temperature recorded was 34.3°C. Over the same period, at the Fowling Point, the minimum temperature was −9.0°C and the maximum temperature was 44.8°C. Salinity ranged from 9.2 PSU to 36.4 PSU. From 2021 to 2023, the lowest temperature recorded in the Wachapreague Channel was 0.002°C and the highest temperature recorded was 33.1°C. Over the same period, at the Upper Haul Over, the minimum temperature was −5.5°C and the maximum temperature was 45.0°C. Salinity ranged from 8.4 PSU to 35.4 PSU.

### Sampling and sex-linked marker efficacy

We identified the reproductive state of 4,855 thalli sampled across all sites and all collection events (Figure 2, Table 1, Table S1). We used the sex-linked PCR assay to determine the phase and for gametophytes, sex of 3,555 thalli, including a subset of thalli with observable reproductive structures at the time of collection. Forty-eight thalli (∼1%) had mismatches between the reproductive state identified at the time of collection or ambiguous bands on agarose gels. Of these mismatches, about half were cystocarpic females at the time of collection but had both female and male bands in the PCR assay.

**Figure 2.**
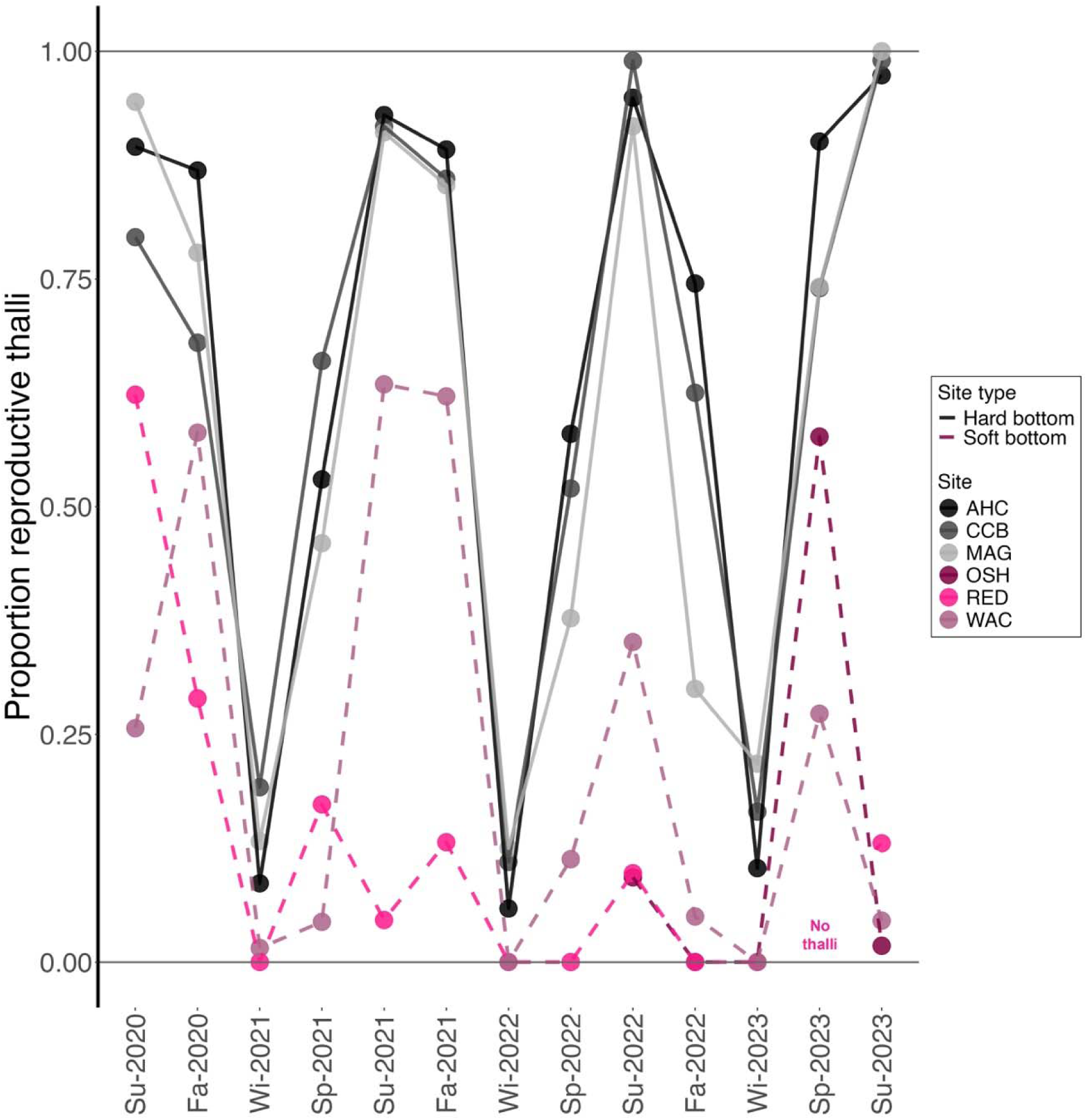
The proportion of reproductive *Gracilaria vermiculophylla* thalli sampled in each season from 2020 to 2023. Hard bottom sites – AHC, CCB, and MAG – are shown as solid lines in shades of black/gray and soft bottom sites – OSH, RED, and WAC – are shown as dashed lines in shades of maroon. OSH was only added as a site in Summer 2022. The proportion of reproductive thalli includes tetrasporophytes, male gametophytes, and female gametophytes.

### Phenology in hard bottom sites

Occasionally, there were free-living thalli found at soft bottom sites. They were rare and we did not sample them. Instead, all thalli sampled from AHC, CCB, and MAG were fixed to hard substrate: at AHC, thalli were fixed to shell hash, old pier pilings, and bricks; at CCB, thalli were all fixed to the jetty of armourstones at the public beach; and at MAG, all thalli were fixed to shell hash and old pier pilings. The average proportion of reproductive thalli was greatest in the boreal summer (∼93% of thalli), between 61% and 73% in spring and fall, respectively, and least in the winter (∼13% of thalli; Table 1, Table S1, Figure 2). The overall patterns of reproductive thalli were similar when only tetrasporophytes (Figure S3a), only gametophytes (Figure S3b), only female gametophytes (Figure S3c), or only male gametophytes (Figure S3d) were considered (note females here were all cystocarpic). Of the sampled gametophytic thalli, the proportion of cystocarpic females and reproductive males together was ∼95% in the summer, 81% in the fall, 26% in the winter, and 54% in the spring. Cystocarpic female gametophytes made up anywhere from 21% to 96% of the female gametophytes sampled across the four seasons. Reproductive male gametophytes made up anywhere from 23% to 93% of the male thalli sampled. In the spring, on average, 80% of sampled males were reproductive as compared to ∼21% cystocarpic females. Tetrasporophytes exhibited greater reproductive periodicity where on average 2% of sampled thalli were reproductive in winter, ∼68% in spring and fall, and 92% in the summer.

Fixed gametophytes composed ∼42% and fixed tetrasporophytes ∼58% of the thalli, though the proportions fluctuated throughout the seasons and across the three hard bottom sites (Figure 3a-c, Figure S4a-c). At each collection time, the *Gracilaria vermiculophylla* populations at AHC, CCB, and MAG were significantly tetrasporophyte biased, except for the first and second winter thallus collections at CCB (Table 1). The *p*-values for the binomial test for a deviation from the expected √2:1 at these two sampling events at CCB were ∼0.07 and therefore close to the nominal significance value of 0.05. Ploidy diversity values ranged from 0.34 to 0.98 (Table 1, Table S1): (i) at AHC, 0.50 < *P_HD_* < 0.88 (mean = 0.68, variance = 0.01); (ii) at CCB, 0.34 < *P_HD_*< 0.98 (mean = 0.75, variance = 0.03); and (iii) at MAG, 0.47 < *P_HD_* < 0.86 (mean = 0.70, variance = 0.01).

**Figure 3.**
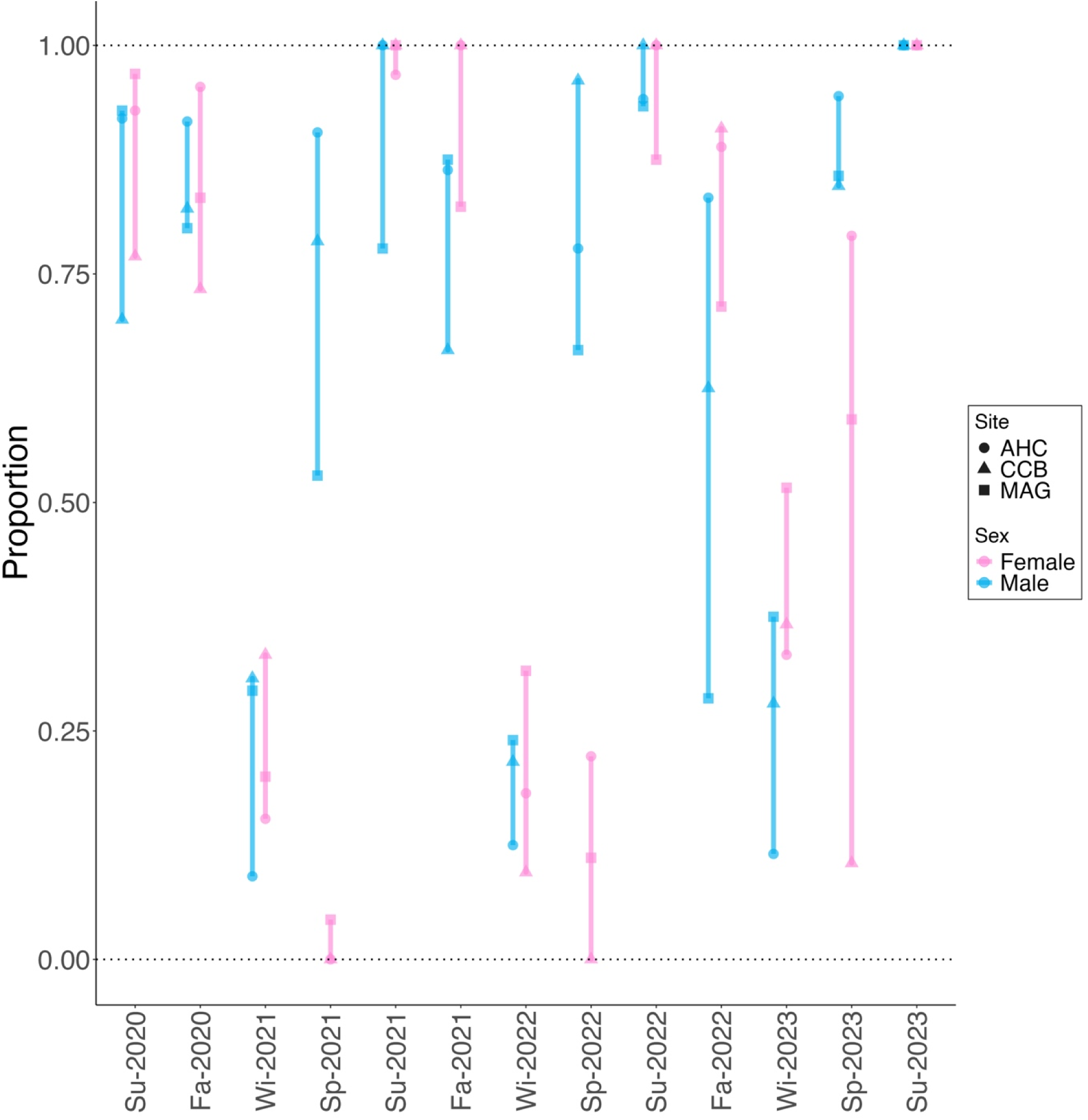
The proportion of reproductive male gametophytes (shown in blue) or cystocarpic female gametophytes (shown in pink) across seasons at the three hard bottom sites – AHC (shown as a circle), CCB (shown as a triangle), and MAG (shown as a square). There is synchrony at all time points except Spring 2021 and Spring 2022 when cystocarpic females lag male gametophytes.

The sex diversity (*P_FM_*) values ranged from 0.45 to 1.00 across all hard bottom sites and sampling events (Table 1, Table S1): (i) at AHC, 0.49 < *P_FM_* < 1.00 (mean = 0.81, variance = 0.02); (ii) at CCB, 0.55 < *P_FM_* < 0.96 (mean = 0.76, variance = 0.02); and (iii) at MAG, 0.45 < *P_FM_* < 0.97 (mean = 0.75, variance = 0.02). Depending on the sampling date, the populations at AHC, CCB, or MAG could be female-(N = 8) or male-biased (N = 9), but many of the observed sex ratios (N = 22) did not differ significantly from a 1:1 ratio at an alpha = 0.05 (Table 1). The adjusted *P_FM_* (by 1×10^−6^) varied significantly by season, with spring having greater *P_FM_* than fall or winter, independent of site or year (Table S3). Differences among hard bottom sites were not observed, though this may be a limitation of sample size (e.g., AHC vs. CCB, *p*-value = 0.09, Table S3a). Spring was associated with greater *P_FM_* compared to fall (Table S2a; Estimate = 1.182, *p* = 0.003), as well as winter (Table S3b; *p* = 0.036). Summer and winter were not different from fall or spring (*p* > 0.150; Table S2). Year-to-year variation was negligible (variance = 1.06 × 10^−10^, standard deviation = 10.03 × 10^−5^). We observed a lag in which males led females by one season (*r*_lag=−1_ = 0.35, permutation test *p* = 0.03), and this appeared to be in the spring in which more males were reproductive than there were cystocarpic females (Figure 3). The cross-correlation coefficient was also significant when there was synchronous cystocarpic females and reproductive males (*r*_lag=0_ = 0.55, permutation test *p* = 0.001).

At each of the hard bottom sites, we observed what appeared to be an inverse relationship between the proportion of reproductive thalli (including tetrasporophytes and gametophytes) and ploidy diversity (*P_HD_*; Figure 4). However, there was only a significant inverse relationship between the proportion of reproductive thalli and *P_HD_* at CCB (Spearman’s ρ = −0.832, *p* = 0.001; Table S4).

**Figure 4.**
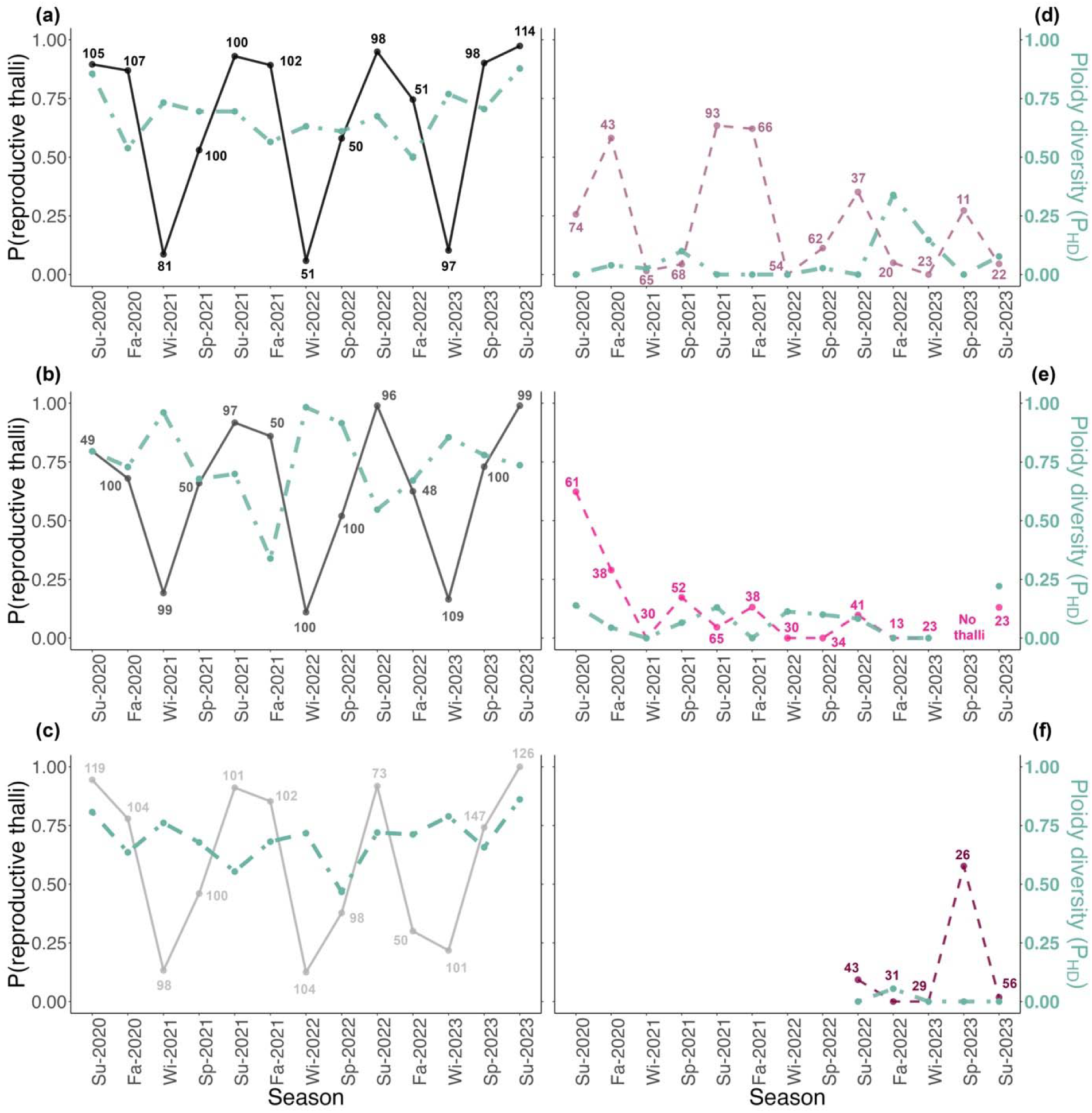
The proportion of reproductive thalli on the left axis (shown as a solid line). On the right axis, is the ploidy diversity (*P_HD_*) for each sampling point (shown as a dashed line). The sample size of thalli is given for each sampling point. a) AHC, b) CCB, c) MAG, d) WAC, e) RED, and f) OSH.

### Phenology in soft bottom sites

By contrast, in soft bottom sites, all thalli were free-living, mostly anchored by the tube building polychaete *Diopatra cuprea*. The number of *Diopatra cuprea* tube caps (Table S5a, as a proxy for the number of polychaetes potentially decorating with *Gracilaria vermiculophylla* thalli, varied among sites and seasons (Table S5b). The variance in tube cap counts was greater than the mean, but the dispersion parameter (θ = 6.74) indicated that overdispersion was adequately handled in the model. The abundance per m^2^ ranged from 13.7 to 36.5 tube caps. At RED, we found significantly fewer *D. cuprea* tube caps as compared to OSH (Estimate = - 0.490, *p* = 0.0003) and WAC (Table S5c, *p* = 0.042). However, there were no differences between the number of tube caps at OSH and WAC (Estimate = - 0.167, *p* = 0.197). There were more *Diopatra cuprea* tube caps in July 2022 (Estimate = 0.421, *p* = 0.016) and February 2023 (Estimate = 0.374, *p* = 0.032) compared to October 2022, though no other seasonal comparisons were significant. The random effect of the quadrat accounted for minor variation across the sampling unit (variance = 0.064).

The proportion of tetrasporophytes, on average, was >96% across all seasons and sites. The proportion of tetrasporophytes was always greater than 80%, with most sites ∼97% tetrasporophytic (Table 1, Table S1). Unlike fixed tetrasporophytes in hard bottom sites, free-living tetrasporophytes in soft bottom sites were never greater than 26%, on average, of thalli collected at a soft bottom site (Figure 4d-f, Figure S4d-f). The maximum proportion of reproductive free-living tetrasporophytes was 63% of the sampled thalli at WAC in summer 2021 (Figure 2, Figure 4, Table S1, Figure S3). Ploidy diversity values ranged from 0.00 to 0.34: (i) at OSH, 0.00 < *P_HD_* < 0.05 (mean = 0.01, variance < 0.001); (ii) at RED, 0.00 < *P_HD_* < 0.22 (mean = 0.07, variance = 0.005); and (iii) at WAC, 0.00 < *P_HD_* < 0.34 (mean = 0.06, variance = 0.009).

At WAC and RED, we observed a less pronounced inverse relationship between the proportion of reproductive thalli (including tetrasporophytes and gametophytes) and ploidy diversity (*P_HD_*; Figure 4), but these relationships were not significant (Table S4). The *P_HD_* was so low at OSH, and we collected thalli for such a short period, further analyses were precluded.

### Phenological differences between hard and soft bottom habitats

The adjusted ploidy diversity (*P_HD_*; by 1×10^−6^) varied significantly by site (Figure 5), in which soft bottom sites with free-living thalli had significantly lower values (*p* < 0.001; Table S6a,b). There was very small variance (variance = 1.51e-09) in the random effect of year. Thus, when populations were sampled in the winter, we observed higher *P_HD_*values (i.e., more gametophytes and tetrasporophytes in hard bottom sites) as compared to the fall (Estimate = 0.648, SE = 0.294, *p* = 0.028), but there were no differences between fall and spring or summer (*p* = 0.264 and *p* = 0.141, respectively), nor any other season pairwise comparisons (Table S6b).

**Figure 5.**
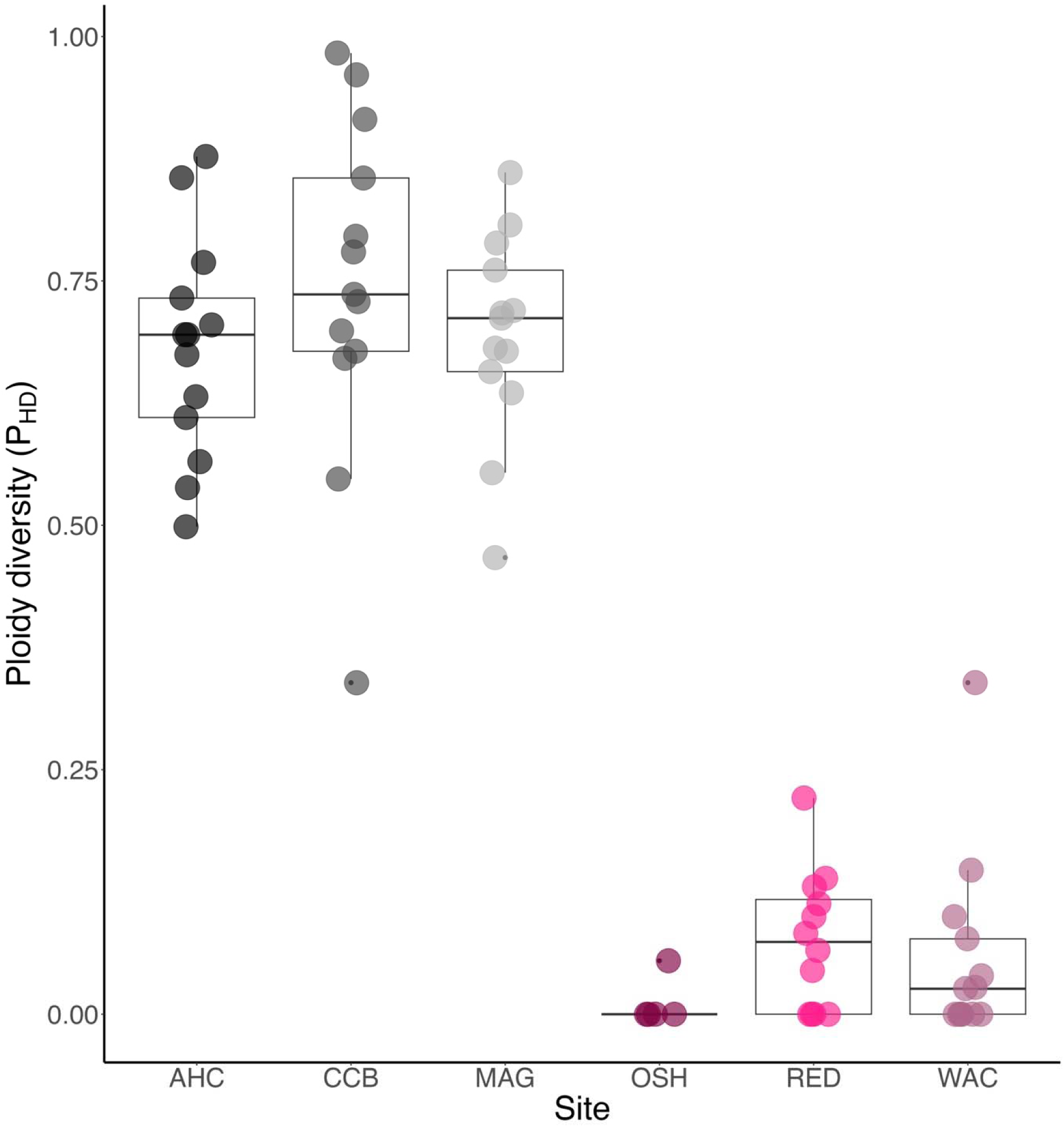
Boxplots showing the distribution of ploidy diversity (*P_HD_*) for hard bottom sites – AHC, CCB, and MAG – and for soft bottom sites – OSH, RED, WAC. Seasonal sampling date is represented as jittered points. The boxes represent the interquartile range with the line showing the median.

Compared to the hard bottom site AHC, the soft bottom at sites OSH, RED, and WAC had strong negative effects on the proportion of reproductive thalli (Table S7a, OSH: Estimate = −3.912, *p* < 0.001; RED: Estimate = −3.407, *p* < 0.001; WAC: Estimate = −2.305, *p* < 0.001), whereas there were no differences between AHC and CCB or MAG (*p* > 0.8). Similar patterns were observed when comparing the other hard bottom sites CCB and MAG to the soft bottom sites (Table S7b). There was very small variance (variance = 5.29 × 10^−10^) in the random effect of year. The greatest proportion of reproductive thalli was found in the summer (Table S7a, Estimate = 1.171, *p* < 0.001; Table S6c, *p* < 0.001). By contrast, winter was associated with a significantly lower proportion of reproductive thalli compared across all seasons (Table S7c, *p* < 0.001). Spring did not, however, differ from fall (Estimate = −0.368, *p* = 0.277).

Finally, temperature was highly associated with the proportion of reproductive thalli, whereby greater temperatures resulted in more reproductive thalli (Table S8a; Estimate – 0.154, *p* < 0.001). Hard bottom sites had much greater proportion of reproductive thalli as compared to soft bottom sites (all comparisons *p* < 0.0001, Table S8a, Table S8b). While there were no differences between hard bottom sites (all *p*-values > 0.99, Table S8a, Table S8b), there were fewer reproductive thalli at OSH as compared to WAC (*p* = 0.035) and thalli at RED were marginally less reproductive than at WAC (*p* = 0.0693).

## DISCUSSION

We surveyed hard and soft bottom habitats in which *Gracilaria vermiculophylla* thalli are now abundant and found strong reproductive periodicity, with a peak in reproductive thalli in the boreal summer, corresponding to the highest temperatures. Hard bottom habitats were composed of both gametophytes and tetrasporophytes. Using a sex-linked PCR assay also allowed us to explore the sex ratio for the first time in this species. We found fluctuations from male- to female-biased across seasons, with the greatest sex diversity (*P_FM_*) in spring in which female and male gametophytes were closer to a 1:1 ratio. However, there was a time lag in which male gametophytes bore spermatangial sori approximately one season earlier than females bore cystocarps. Regardless of substrate type, there was tetrasporophytic bias across sites, characteristic of this genus (see also Kain & Destombe, 1995) and supporting earlier work contrasting native and non-native populations in *G. vermiculophylla* (Krueger-Hadfield et al., 2016). Tetrasporophytic bias was even more pronounced in the soft bottom sites OSH, RED, and WAC in which gametophytes were largely or entirely absent. The dominance of tetrasporophytes was stable throughout the sampling period and across sites. Moreover, soft bottom habitats with free-living thalli had significantly lower proportions of reproductive thalli, even in the peak reproductive period in the summer. Surprisingly, there appeared to be an unexpected inverse relationship between the proportion of reproductive thalli and ploidy diversity (*P_HD_*), though it was only significantly so at the hard bottom site CCB. Below, we discuss the phenology of *G. vermiculophylla* in hard and soft bottom habitats.

### Phase dominance

Phase dominance is well documented across algae, in which populations may be dominated by sporophytes or gametophytes (Hawkes, 1990; see also Krueger-Hadfield, 2020). For example, Dyck and DeWreede (1995) found a dominance of gametophytes in the summer and tetrasporophytes in the winter in the red alga *Mazzaella splendens*, though tetrasporophytes maintained more stable densities throughout the year. Sexual reproduction likely predominated in this population of *M. splendens* with isomorphic gametophytes and tetrasporophytes adopting different strategies leading to ecotypic variation (see also Hughes & Otto, 1999). Yet, the phycological literature is replete with examples of ‘direct life histories’, in which only sporophytes or only gametophytes are found in natural populations or following culturing in laboratory conditions from field collected thalli (e.g., Maggs, 1988). Yet, while we appreciate algal life cycle diversity, we understand less about the spatial patterning of these variants and their corresponding reproductive modes (Krueger-Hadfield, 2020; 2024).

Patterns of geographic parthenogenesis in which there is spatial patterning of sexual and asexual populations are common across taxa (e.g., Haag & Ebert, 2004). Often for macroalgae, these patterns result in sporophytic dominance. For example, Maggs (1998) documented tetrasporophyte-only populations of the red alga *Dasya ocellata* in Ireland and England as compared to populations with alternating gametophytes and tetrasporophytes in Morocco. Not all patterns of asexual reproduction and the dominance of one phase are associated with latitude. Hwang et al. (2005) found differences in *Dictyota spathulata* (as *D. dichotoma* in the original paper, but see Vieira et al., 2025) from the coastline of Korea where populations from the east coast are annuals and asexual. Asexual reproduction leads to sporophyte-only populations, recycling the sporophytic phase via monospores. Sexual populations of *D. spathulata* are found in some localities on the south coast and exclusively on the west coast of the Korean Peninsula, suggesting current patterns and seawater temperature may play a role in the distribution of sexual and asexual populations. Similarly, Gabrielson et al. (2002) found the increasing prevalence of asexual reproduction via paraspores or vegetative propagation in the Kattegat and Baltic Sea, respectively, in *Ceramium tenuicorne* as compared to the Oslofjorden, where gametophytes and tetrasporophytes were common, suggesting sexual reproduction. The gradient in salinity from the Oslofjorden to the Baltic Sea likely favors asexual reproductive modes. Thus, many environmental conditions can drive the spatial patterning of sexual and asexual reproduction in macroalgal populations, though this requires more spatially explicit sampling combined with appropriate genetic tools (e.g., sex-linked markers to assign phase/sex and microsatellites to determine sexual and asexual rates).

Unsurprisingly, in the Mid-Atlantic region where *Gracilaria vermiculophylla* is now a dominant species, tetrasporophytes dominated throughout the seasons and across sites. Previous work had compared native to non-native populations, in which most native thalli were collected from hard bottom habitats and many non-native thalli were collected from soft bottom habitats (e.g., Krueger-Hadfield et al., 2016). This led to an assumption that the invasion of the Northern Hemisphere resulted in the breakdown of the haploid-diploid life cycle, in which tetrasporophytes came to dominate populations. Sample sizes from Japanese populations precluded robust assessments of phase ratios (see sample size discussion in Krueger-Hadfield & Hoban, 2016), but gametophytes and tetrasporophytes were common at all sites sampled (Krueger-Hadfield et al., 2016; 2017). The exceptions were two sites with soft bottom in which only tetrasporophytes were found: one site, Futtsu, was sampled and included in studies investigating the invasion history (e.g., Krueger-Hadfield et al., 2017; Sotka et al., 2018) and another site in Hokkaido was not sampled extensively or included in downstream assays (S.A. Krueger-Hadfield, *personal observations*). Subsequent sampling along the western coast of North America revealed many hard bottom habitats with gametophytes and tetrasporophytes, whereas all soft bottom sites were tetrasporophyte-dominated (Krueger-Hadfield et al., 2018). Together these past observations and our current survey in the Delmarva are strongly suggestive that the substrate type leads to tetrasporophytic dominance, though the mechanisms that lead to the loss of the gametophytes remain enigmatic (see Krueger-Hadfield & Ryan, 2020; Lees et al., 2018).

Previous work in *Gracilaria vermiculophylla* had to rely on reproductive structures and the multilocus genotype from microsatellite loci to document tetrasporophytic dominance (e.g., Flanagan et al., 2021; Gerstenmaier et al., 2016; Krueger-Hadfield et al., 2016; 2017). Rueness (2005) documented the preponderance of vegetative thalli in French collections of *G. vermiculophylla*, rendering studies challenging as phase could not be distinguished. Yet with sex-linked markers (Krueger-Hadfield et al., 2021), we can easily document the phase and sex, if applicable, of all thalli collected from a site (see also Krueger-Hadfield et al., 2023). Tetrasporophytes dominate all sites in both hard and soft bottom habitats, and this pattern was stable from 2020 to 2023. Other studies have found stability in phases ratios through time (see as an example, Thornber & Gaines, 2003). A tetrasporophyte bias may hint that tetrasporophytes are more fit, especially when conditions deteriorate. Experimental manipulations of tetrasporophytes and gametophytes are required to test this hypothesis.

It is curious that ploidy diversity (*P_HD_*) fluctuates, seemingly inversely with the proportion of reproductive thalli. This pattern was only significant at CCB and harder to visualize at each of the soft bottom sites where *P_HD_* values were consistently low. *Gracilaria vermiculophylla* thalli at CCB are fixed to armourstones that create the jetties at the public beach, a much more stable substrate than the thalli at AHC or MAG found more often on shell hash. At AHC, as an example, we have frequently observed thalli fixed to small pieces of shell hash undulating with water movement at low tide (S.A. Krueger-Hadfield & W.H. Ryan, *personal observations*). At CCB, in two winter sample collections, the phase ratio was not distinct from the predicted ratio of √2:1. Thus, the thalli at CCB may be most like the rocky shore populations more often studied in other macroalgae, such as *Mazzaella splendens* (Dyck & DeWreede, 1995) in which there are seasonal patterns of phase dominance or reproduction. In winter in hard bottom habitats, there were more gametophytes as compared to summer, during the peak period of reproduction. It is possible that tetrasporophytes are longer-lived than the gametophytes. Gametophytes did tend to survive at lower rates than tetrasporophytes when grown under laboratory conditions (Krueger-Hadfield & Ryan, 2020). This would set *G. vermiculophylla* apart from its congeners *G. gracilis*, in which there were no differences among phases/sexes in survivorship (Engel et al., 2001), and *G. chilensis* in which female gametophytes had the highest survival (Vieira et al., 2018). Instead, *G. vermiculophylla* would be more like *M. splendens*, in which Dyck and DeWreede (1995) found lower survivorship and shorter life spans for gametophytic ramets (physically separate module of a genet, e.g., a frond). This indeed raises a salient point that often these studies are working on the ramets, rather than genets (a single genetic individual, e.g., a holdfast and all its fronds). Engel et al. (2001) and Vieira et al. (2018) did track genets, but we sampled a single, terete upright and therefore are documenting ramet dynamics in *G. vermiculophylla*. In the Mid-Atlantic, there is strong seasonality that may influence gametophytic vs. tetrasporophytic survival and may be a fruitful avenue worth pursuing to understand tetrasporophytic dominance in general but also specifically following the invasion of soft bottom habitats.

### Fluctuations in male or female gametophytic proportions in hard bottom habitats

Few studies of macroalgal phenology have been able to differentiate sexes when thalli are not reproductive (e.g., Dyck and DeWreede, 1995; Kamiya et al., 2021; Krueger-Hadfield et al., 2013; Thornber & Gaines, 2003). Exceptions include Vergés et al. (2008) in *Asparagopsis armata*, in which the sex ratio was 1:1 at the beginning of the reproductive season, becoming female biased by the end of the season in Australia. Further, Mihaila et al. (2023) found spermatangia were present on gametophytic thalli in August and September with the appearance of cystocarps occurring in September in New Zealand. Observational data still relies on the presence of reproductive structures. The use of sex-linked markers has only been applied to field surveys in a few taxa as few macroalgae have these types of markers identified. Data on sex ratios were not reported by Couceiro et al. (2015) in the sexual populations of *Ectocarpus siliculosus* and *E. crouaniorum*, even though sex-linked markers were used to assign phase. Only 16 gametophytes were found out of 182 specimens assayed with sex-linked markers by Arai et al. (2024), precluding robust analyses of sex ratios across populations in *Dictyota dichotoma* (note this likely *D. spathulata,* see Vieira et al., 2025). In *G. chilensis*, sex linked markers were not used to explore sex ratios but were used instead to model demographic parameters in which there were no differences found between phases or sexes when not reproductive (Vieira et al., 2018). Instead, females grow faster and larger than tetrasporophytes and males (Vieira et al.; 2021) and ultimately survive better (Vieira et al., 2018).

In *Gracilaria vermiculophylla*, Terada et al. (2000) found reproductive male gametophytes were present in June before cystocarpic female gametophytes were observed later in July at a site studied in Hokkaido, Japan. Similarly, Muangmai et al. (2014) found reproductive male gametophytes in November before observing cystocarpic female gametophytes in December onward. Nevertheless, vegetative thalli dominated the study site in Terada et al. (2000) and made up ∼10-15% of the thalli in Muangmai et al. (2014). In our study, we were able to determine the phase of every thallus we sampled, regardless of reproductive state. We observed a lag in spring in which reproductive males were more abundant than cystocarpic females. In *G. gracilis,* cystocarps reach maturity about one month after fertilization (Engel et al., 2001). Reproductive females were no doubt present in these *G. vermiculophylla* populations, including at CCB where there were no cystocarpic females sampled in April 2021 and April 2022. Following fertilization, carposporophytes develop and cystocarpic females increased in frequency. Future studies should occur more regularly to capture immature cystocarps and obtain a more fine-tuned phenological calendar.

We also found fluctuations across seasons in the sex ratio for *Graclaria vermiculophylla* gametophytes at AHC, CCB, and MAG, though most observations were not significantly different from a 1:1 ratio. Krueger-Hadfield et al. (2021) showed that there is likely Mendelian segregation of the sex determining region, further explored in Lipinska et al. (2024), suggesting there should be a 1:1 sex ratio in populations that complete the haploid-diploid life cycle – in other words, in hard bottom habitats. Yet, our observation of greater sex diversity (*P_FM_*) in spring, indicating a sex ratio closer to 1:1, with variation in the proportions of females and males across space and seasons, warrants further investigation. For example, under nutrient-rich conditions *G. vermiculophylla* male gametophytes showed accelerated growth while females did not from collections at one of the same sites we surveyed here, MAG (Krueger-Hadfield & Ryan, 2020). Similarly, there were hints of differences between female and male gametophytes in biomechanical properties and thallus composition at a single site in South Carolina, but these patterns were swamped by inter-annual variation (Lees et al., 2018). It is plausible that some of these fluctuations may reflect sampling stochasticity. To tease apart the phenotypic differences between males and females from sampling artifacts, further experimental approaches are needed.

### Diopatra cuprea *and* Gracilaria vermiculophylla *in soft bottom habitats*

In regions where there are species of the tube building polychaete genus *Diopatra*, there is an association between the polychaete and *Gracilaria vermiculophylla* (see as examples, Abreu et al., 2011; Kollars et al., 2016; Mott et al., 2022; 2025; Thomsen & McGlathery, 2005). Most of the biomass collected in 0.25 m^2^ quadrats at the soft bottom sites OSH, RED, and WAC in our study were anchored to the tube caps of *D. cuprea*. Worm abundance fluctuates over seasons and likely also plays a role in *G. vermiculophylla* biomass by anchoring thalli in the photic zone. We found more *D. cuprea* in the summer as compared to the other months surveyed from 2022 to 2023. We also observed differences across the three soft bottom sites in which there were fewer *D. cuprea* at RED as compared to OSH or WAC. Curiously, there were no *G. vermiculophylla* thalli at RED in Spring 2023, and subsequent visits to this site also revealed no thalli in one visit in spring 2024 and one in winter 2025 (S.A. Krueger-Hadfield, *personal observations*). While there were no differences in *D. cuprea* abundance between OSH and WAC, *G. vermiculophylla* thalli also were not present during an unrelated visit to OSH in winter 2025 (S.A. Krueger-Hadfield, *personal observations*). By contrast, though the biomass fluctuates, the *G. vermiculophylla* population at WAC seems the most stable of the three soft bottoms sites studied (see Krueger-Hadfield & Ross, 2025). Keller et al. (2019) documented evidence of *D. cuprea* declines in regions where *G. vermiculophylla* is now found. Moreover, the authors found elevated *D. cuprea* mortality at WAC (called Worm Flat, Burton’s Bay in their study) associated with exceptionally high biomass of *G. vermiculophylla*. However, *G. vermiculophylla* has been in these environments for much longer than appreciated in the literature and documented by Keller et al. (2019), suggesting the influence of the *G. vermiculophylla* on *D. cuprea* is more complex. We sampled the same site as Berke (2012) and found slightly lower average numbers of *D.* cuprea per m^2^ (Berke, 2012: 36.9 worm/m^2^; our study from the two summer collections: 30 worms/m^2^). However, regular surveys are required to determine the magnitude of inter-annual variation. We do not have quantitative data on the biomass of *G. vermiculophylla* that exists in the intertidal versus subtidal zones in the coastal bays along the Delmarva. Since *D. cuprea* facilitates algal distributions in Virginian coastal bays (Thomsen & McGlathery, 2005) and actively chooses *G. vermiculophylla* over other decoration options (Mott et al., 2022), the links between the stability of *G. vermiculophylla* populations to the *D. cuprea* abundance are curious and worthy of further investigation.

### Marine macroalgae in Virginian coastal bays

The prevalence of sandy beaches and mudflats in the Mid-Atlantic region is “without any doubt, the reason why no phycologist has ever focused [their] attention to this interesting area and why this area still forms a gap in our knowledge of the algal distribution along the East Coast of the United States” (pg. 17, Zaneveld & Barnes, 1965). Humm (1979) further noted that the knowledge of the benthic marine macroalgae in Virginia were not well known, with surveys focused northward on New England and southward to the Carolinas. Indeed, Mathieson and Dawes (2017) and Schnieder and Searles (1991) leave a gap along the Delmarva (see also Kapraun, 1980). The last surveys were undertaken by Orris (1980) and Connor (1980) from surveys in the early to mid-1970’s in the Chesapeake Bay in Maryland, with only one site (#15 in Connor, 1980 near our site AHC). Thus, our knowledge of distributions is both geographically and temporally limited, with surveys nearly 50 years old. Nevertheless, studies in this area documented seasonal floras corresponding to higher and lower latitude species that reach their limits in this region.

The reproductive periodicity was documented for a variety of red, green, and brown macroalgae by Zaneveld and Barnes (1965), including thalli that could plausibly be *Gracilaria vermiculophylla*. The authors note two species of *Gracilaria*: (i) *G. foliifera* as a component of the fall and winter floras; and (ii) *G. verrucosa* – a taxonomic catch-all for gracilarioid taxa – is a perennial. Moreover, Rhodes (1970) also notes that *G. foliifera* is rare in winter and common in summer, whereas *G. verrucosa* is abundant in the summer. *Gracilaria foliifera* was later determined to be *G. tikvahiae* (Edelstein et al., 1978; McLachlan, 1979), a species with flattened thalli as compared to the terete thalli of other gracilarioid thalli, such as in Humm (1979). These *G. verrucosa* thalli strongly resemble *G. vermiculophylla*. Figure 31 (pg. 26) in Zaneveld and Barnes (1965) shows *Gracilaria* thalli growing on a boulder at Fort Wool on the western shore near the Hampton Roads area at the mouth of the James River in Virginia. These thalli look very similar to the habit of *G. vermiculophylla* thalli at CCB, growing luxuriantly on armourstones. Thomsen et al. (2005) sequenced a thallus collected in 1998 from three localities in Hog Island Bay, including Oyster Harbor, a site included in Krueger-Hadfield et al. (2016; 2017) and Mott et al. (2022) and near the site OSH in our current study. The thallus was *G. vermiculophylla* which has led to the conclusion that *G. vermiculophylla* arrived in Virginia in 1998 (Thomsen et al., 2005, see also Keller et al., 2018). However, it is likely much earlier based on the history of moving oysters, a main vector for transporting *G. vermiculophylla* thalli across ocean basins (Krueger-Hadfield et al., 2017), and records of terete thalli in the coastal bays of the Delmarva. This underscores the necessity of monitoring the seasonal patterns of macroalgae.

### The evolutionary ecology of free-living algae

The consequences of seasonal variation in temperature, photoperiod, irradiance quantity, and nutrient levels have been documented for macroalgae (e.g., Murray & Dixon, 1992). de Bettignies et al. (2018) found that temperature controls the maturation of spores and gametes in a review of 81 studies. Yet, Norton and Matheison (1983) concluded free-living algae are sterile and do not become reproductive, though most of their review was on fucoid ecads. Krueger-Hadfield et al. (2023) demonstrated free-living *Gracilaria vermiculophylla* tetrasporophytes do become reproductive and peak in the summer with rising temperatures. However, the authors sampled a single soft bottom site in which the peak in the proportion of reproductive thalli was 65%. In our study, we observed differences between hard and soft bottom sites in the proportion of reproductive thalli. The maximum proportion of reproductive thalli was 63% in soft bottom habitats, with an average of 23% across summer sample collections. Variation was also considerable across soft bottom sites in the summer within and among OSH, RED, and WAC, with as few at 2% of the tetrasporophytes were reproductive. We found more reproductive tetrasporophytes at WAC, a site that seems to be more stable in terms of *G. vermiculophylla* biomass (Krueger-Hadfield & Ross, 2025). While temperature may be a significant predictor of an increase in reproductive thalli (gametogenesis and sporogenesis, see also de Bettignies et al., 2018), it does not explain the stochasticity across soft bottom sites, nor the differences between hard and soft bottom habitats. Guillemin et al. (2013) demonstrated that vegetative tetrasporophytes grew faster than reproductive thalli. If similar patterns occur in *G. vermiculophylla*, this could explain the dominance of vegetative thalli across soft bottom sites, further facilitated by breakage of thalli by *Diopatra cuprea*. The amount of site-specific variation in soft bottom habitats suggests lumping sites under the categorical variable of soft bottom is likely to obscure variability among habitats.

While this study has begun to fill in some of the gaps outlined as future work by Norton and Mathieson (1983), there remain many enigmatic features of fixed and free-living thalli. Krueger-Hadfield et al. (2023) found mixed phase thalli in which cystocarpic females also bore tetrasporangial sori. At the time of collection, we found cystocarpic females that subsequently had two bands following the sex-linked PCR assay, resembling tetrasporophytes. Cystocarps bear both maternal and paternal DNA (see Krueger-Hadfield et al., 2015), possibly generating female and male bands if cystocarps were included in the DNA extraction. Alternatively, if thalli are mixed phase, as found in other *Gracilaria* species (Kain & Destombe, 1995), then the presence of male and female bands in the sex-linked PCR assay may reflect the presence of tetrasporophytic and female gametophytic regions of the thallus (see van der Meer, 1986 as an example). We did not have herbarium mounts for all thalli that had discrepancies between the field identification and the sex-linked phase determination. For those with herbarium mounts, we did not observe any tetrasporangial sori (S.A. Krueger-Hadfield, *personal observations*). In addition, thalli were completely extirpated from RED in spring 2023, but thalli were present in the next sampling period in summer 2023. The connection between and among hard and soft bottom populations is unknown, though patterns of genetic differentiation were lower when performing pairwise comparisons between Delmarva sites in contrast to sites sampled in New England or the southeastern United States (Krueger-Hadfield et al., 2017). Finally, discovering the ecological factors that contribute to tetrasporophyte bias in *G. vermiculophylla* in both hard and soft bottom populations is a critical next step. These data will not only contribute to our understanding of the expansion of *G. vermiculophylla* from its native range but also of the cycling between ploidy phases which will lead to a broader understanding of life cycle diversity and the evolutionary ecology of sex.

## ACKNOWLEDGEMENTS

S.A. Krueger-Hadfield dedicates this manuscript to the memory of Dr. Leonard J. Dyck.

We thank S. Fate, P.G. Ross, R. Snyder, E. Smith, J. Lewis, H.F. Parks, G. Brundage, and J. Paul for logistical support at the Virginia Institute of Marine Science Eastern Shore Laboratory (VIMS ESL). We also thank R. Hadfield, B. Thornton, T. Williams, A. Brinegar, O. Melendez Vera, and H. Rippin for additional field and lab assistance. The UAB Department of Biology provided logistical support for travel to VIMS ESL. Support for this project was provided by start-up funds from the College of Arts and Sciences at The University of Alabama at Birmingham (to SAKH), the International Phycological Society Paul Silva Student Grant (to APO), the National Science Foundation CAREER Award (DEB-2141971 [UAB] and DEB-2436117 [VIMS|WM] to SAKH), and NSF OIA-1946412 (to SAKH). The water quality monitoring station at Wachapreague was supported with funds from R.A. Snyder at VIMS ESL and the station at Willis Wharf was supported by donations from S. and B. Johnsen. We also thank Cherrystone Aquafarms for providing the site support and utilities for the Willis Wharf station. The Owens Foundation at VIMS ESL partially supported A. Brinegar during her visit at VIMS ESL. H. Rippin and O. Melendez Vera were supported by the Bonnie Sue Summer Scholarship. SAKH was supported by the NSF CAREER award (DEB-2141971), NSF EAGER award (DEB-2113745), and the Norma Lang Early Career Fellowship from the Phycological Society of America. APO was supported by a UAB Blazer Fellowship. WHR was supported by the NIH-IRACDA MERIT Post-doctoral Fellowship program at UAB (NIH K12GM088010).

## AUTHOR CONTRIBUTIONS

**Stacy A. Krueger-Hadfield** Conceptualization (lead), Data curation (lead), Formal analysis (lead), Funding acquisition (lead), Investigation (lead), Methodology (lead), Project administration (lead), Resources (lead), Supervision (lead), Visualization (lead), Writing – original draft (lead), Writing – review and editing (equal); **Alexis P. Oetterer** Data curation (equal), Investigation (supporting), Writing – review and editing (equal); **Darian M. Kelley** Data curation (equal), Investigation (supporting), Methodology (supporting), Resources (supporting), Writing – review and editing (equal); **Will H. Ryan** Conceptualization (supporting), Investigation (supporting), Methodology (supporting), Writing – review and editing (equal)

## DATA ACCESSIBILITY

Data and code are provided at Zenodo (TBD).

## ABBREVIATIONS

AHC: Ape Hole Creek
bp: base pair
BSA: bovine serum albumin
CCB: Cape Charles Beach
MAG: Magotha Rd.
OSH: Hillcrest
PCR: polymerase chain reaction
PPT: parts per thousand (salinity)
RED: Fowling Point
VIMS ESL: Virginia Institute of Marine Science Eastern Shore Laboratory
WAC: Upper Haul Over

---

**Figure S1.**
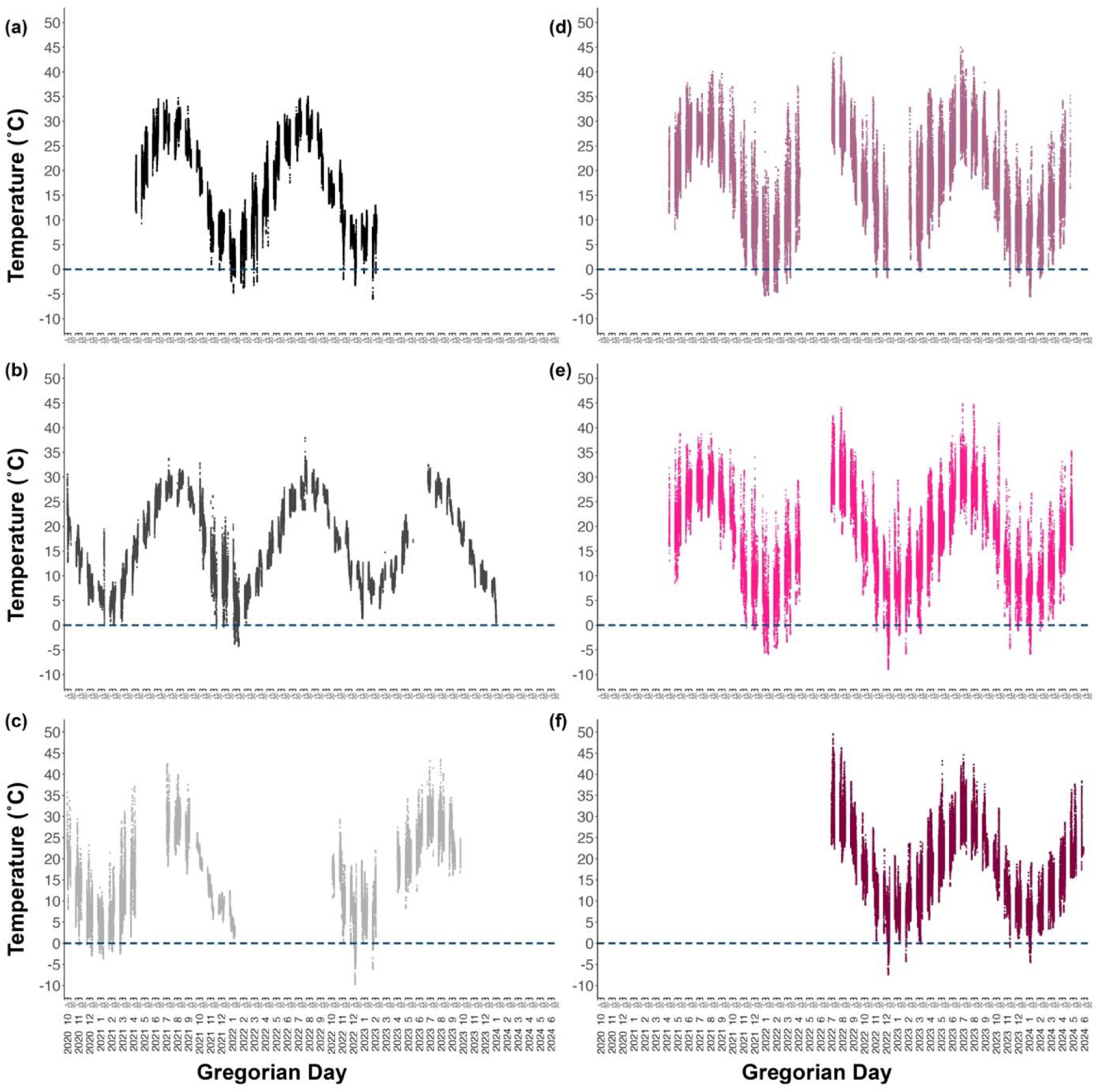
Temperature profiles for each site with Gregorian Day shown on the x-axis from HOBO loggers deployed at each site (see Methods). The year and month are provided along the x-axis with 1, 15, 30 to span each month. Points are shown from multiple measurements for each Gregorian day. a) AHC, b) CCB, c) MAG, d) WAC, e) RED, and f) OSH.

**Figure S2.**
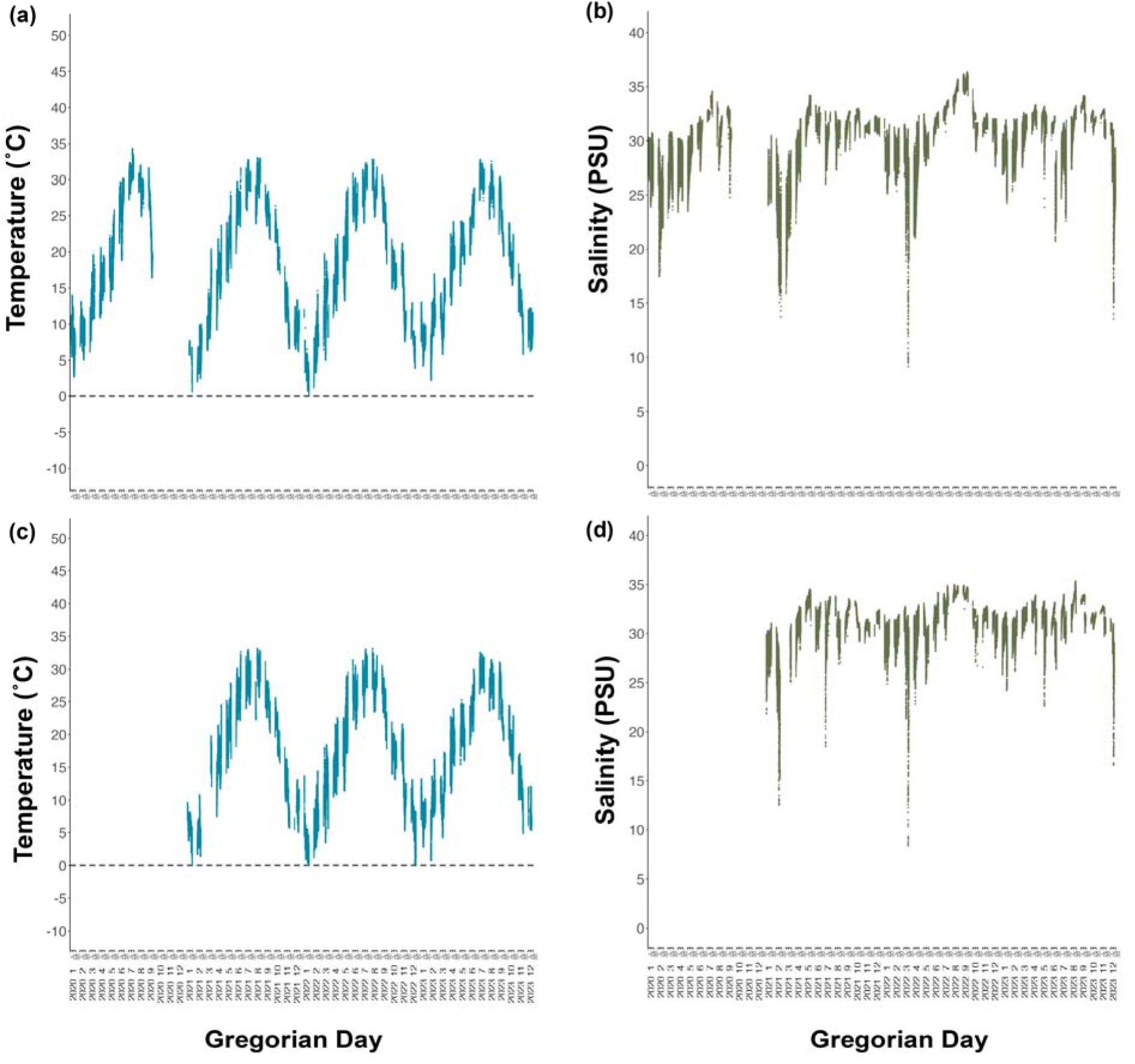
Temperature (°C) and salinity (PSU) plots from the a) and b) Willis Wharf station and c) and d) Wachapreague station. Temperature and salinity profiles for each site with Gregorian Day shown on the x-axis with the year and month are provided with 1, 15, 30 to span each month. Points are shown from multiple measurements for each Gregorian day.

**Figure S3.**
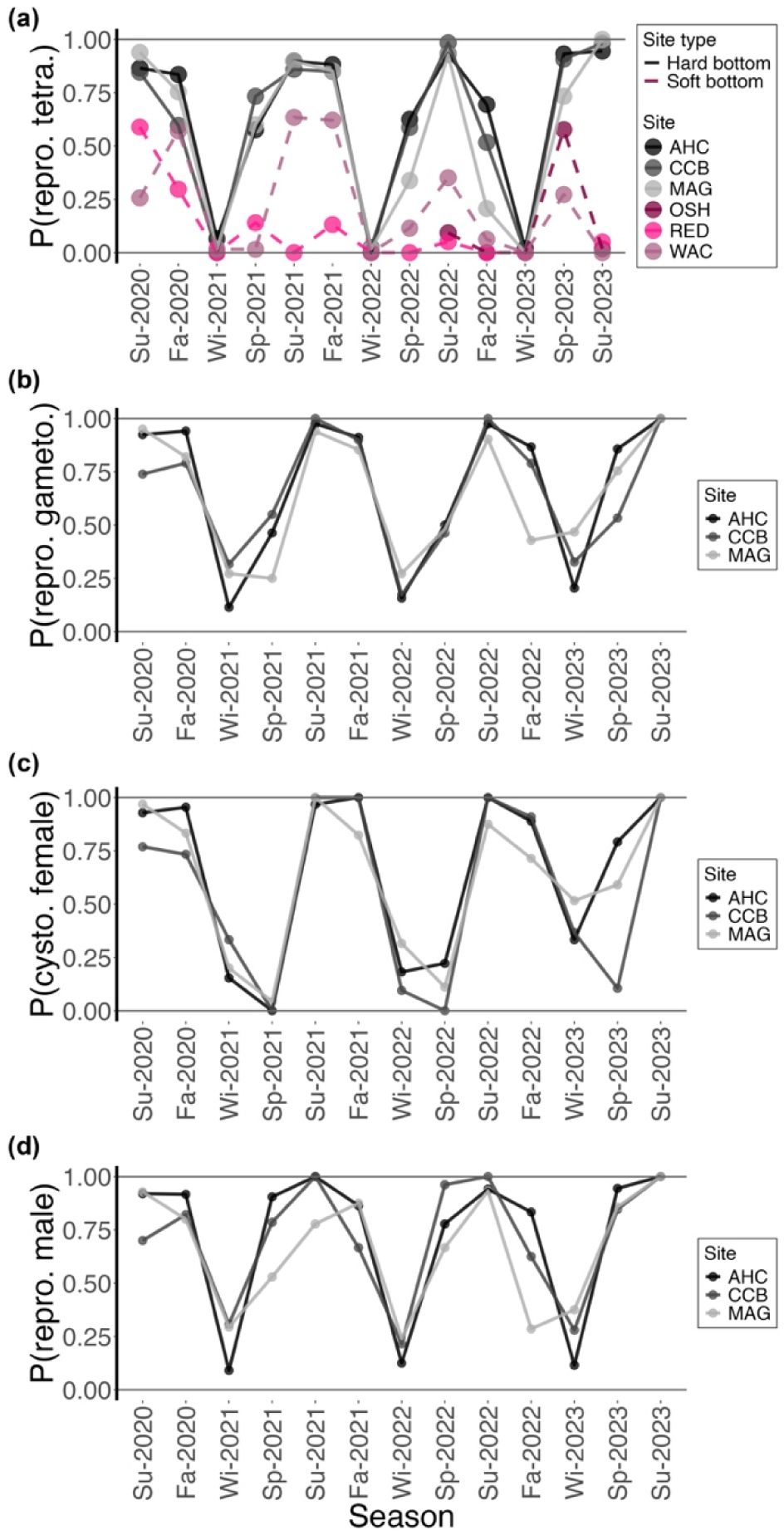
The proportion of reproductive *Gracilaria vermiculophylla* thalli sampled in each season from 2020 to 2023. Hard bottom sites – AHC, CCB, and MAG – are shown as solid lines in shades of black/gray and soft bottom sites – OSH, RED, and WAC – are shown as dashed lines in shades of maroon. OSH was only added as a site in Summer 2022. a) The proportion of reproductive tetrasporophytes, b) proportion of reproductive gametophytes (including both cystocarpic females and reproductive males), c) the proportion of cystocarpic females, and d) the proportion of reproductive males.

**Figure S4.**
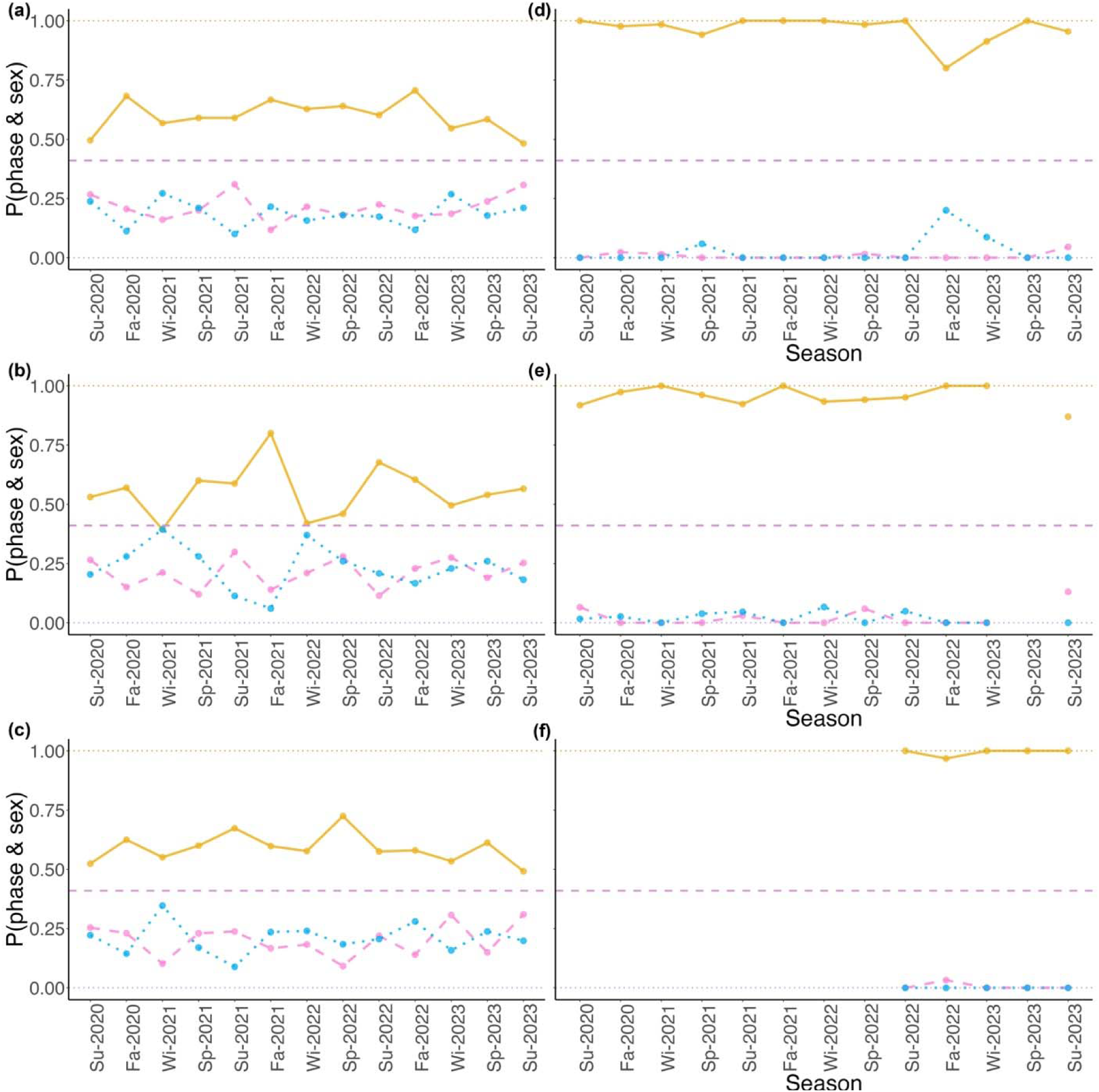
The proportion of tetrasporophytes (shown in goldenrod and a solid line), female gametophytes (shown in pink and a dashed line), and male gametophytes (shown in blue and a dotted line) for a) AHC, b) CCB, c) MAG, d) WAC, e) RED, and f) OSH. The lavender line at0.41 shows the expected proportion of √2:1.

## Notes

### Competing Interest Statement

The authors have declared no competing interest.

